# Inhalable Perfluorocarbon RNA Nanocapsules Bypass Immune Clearance While Targeting Lung Epithelial and Lung Tumor Cells

**DOI:** 10.1101/2025.06.05.658088

**Authors:** Kasturi Siddhanta, Atefehsadat Monirvaghefi, Aditya Sundar, Braeden R. Pinkerton, Neha Kumari, Ling Ding, C. J. Woslager, Marjina Akter Kalpana, Chinmay M. Jogdeo, Ashley R. Ravnholdt, Jill A. Poole, Joshua L. Santarpia, James E. Talmadge, David Oupický

## Abstract

Inhalation RNA therapy offers to transform treatment of pulmonary diseases, yet mucus trapping, immune clearance, and navigation of heterogeneous lung tissue architecture still prevents RNA from reaching its target cells. Here, we develop perfluorocarbon (PFC) RNA nanocapsules that show negligible immune clearance, minimal inflammatory response, and efficient mucus transport, while passively homing to lung epithelial and tumor cells. After a single aerosolized dose in orthotopic lung metastasis model, more than 60% of tumor cells and most type II alveolar and bronchial epithelial cells internalized the nanocapsules, with observed pulmonary retention exceeding 48 h. The nanocapsule provoke negligible cytokine release, enabling repeated dosing. Treatment with therapeutic miR34-a suppresses metastatic outgrowth, potentiates anti-tumor immunity, and almost doubles median survival relative to control paclitaxel chemotherapy. By combining unique PFC disposition with RNA versatility, the delivery platform overcomes the main biological barriers for inhalable RNA medicines and opens a translatable path for treating diverse pulmonary diseases.

## Introduction

Pulmonary delivery of therapeutic RNA via inhalation offers a promising strategy for treating lung diseases, including primary lung tumors and metastases ^1^. This approach provides direct access to the airways, enabling localized treatment and high drug concentrations at the site of action. Consequently, this method facilitates rapid clinical responses while minimizing systemic exposure and associated systemic adverse effects ^2^. By circumventing first-pass metabolism, inhaled therapeutics may achieve equivalent or greater efficacy compared to systemic administration at a fraction of the dose ^3^. Furthermore, inhalation is a non-invasive route of administration, enhancing patient comfort and compliance ^1,2^. Recent advances in lipid nanoparticle (LNP) and aerosolization technology have further propelled the potential of inhaled RNA therapeutics to effectively target and treat lung metastases ^4–6^. Furthermore, the development of novel microfluidic platforms has enabled shear-free generation of LNP aerosols, preserving their structural integrity and improving delivery efficiency ^7^. Despite the pre-clinical advances in inhalable pulmonary delivery systems, clinical translation remains in early stages due to challenges such as rapid immune clearance and observed toxicity ^8,9^.

Designing successful pulmonary delivery systems requires careful optimization of particle properties, including size, morphology, surface characteristics, cell targeting and evasion of host defenses ^3^. The deposition of inhaled particles in the respiratory tract is governed by the aerodynamic diameter of the aerosol particles which represents the diameter of a unit-density spherical particle (1 g/cm³) that has the same settling velocity in air as the particle being measured. Particles of 1-5 μm being optimal for lower airway targeting, while particles > 6 μm deposit in upper airways and those < 0.5 μm risk being exhaled ^10^. The presence of lung tumors can obstruct the airways and divert airflow patterns, potentially hindering microparticle delivery to the tumor site ^11^. Simulations indicate that particles accumulate increasingly at tumor sites until the tumor growth obstructs roughly 50% of the airway passage, after which particle deposition diminishes as blockage becomes more severe ^12^. Therefore, to effectively target lung tumors, both the nanoparticle formulation and its aerosolized form must be optimized. The aerosol delivery parameters, including particle aerodynamic diameter, density, shape, and dispersibility, govern respiratory tract deposition patterns and determine whether particles reach the targeted lung regions. Meanwhile, the nanoparticle formulation itself must facilitate retention within the targeted lung tissue, enhance permeability into tumors, and evade clearance mechanisms like mucociliary transport and phagocytosis ^13^. This can be achieved through various particle engineering strategies. For dry powder formulations, creating inhalable aggregates, where nanoparticles are co-formulated with other agents to form particles with aerodynamic diameters of 1-5 μm, helps overcome the tendency of individual nanoparticles <500 nm to agglomerate due to strong cohesive forces and poor dispersibility. For liquid formulations, strategies include nebulization of nanoparticle suspensions with appropriate surfactants to maintain stability and prevent aggregation during aerosolization. Regardless of formulation type, other approaches include surface modification by surfactant coating, active targeting using tumor-specific ligands, reducing phagocytic clearance using large porous particles, or using anti-fibrotic agents that can decrease tumor interstitial fibrosis and improve the intratumoral distribution of nanoparticles ^11^.

Perfluorocarbons (PFCs) offer multiple benefits for pulmonary RNA therapy through several biophysical and biological mechanisms that enhance the delivery, distribution, and overall transfection efficiency. Their unique properties, including high density and low surface tension at PFC-air interfaces enable them to readily spread and form thin films over the hydrophobic components of the lung surfactant, thus facilitating improved pulmonary distribution of delivery vectors by potentially penetrating mucus and ensuring uniform spread across the alveolar surface ^14–16^. For instance, combined pulmonary delivery of perflubron and adenoviral vectors resulted in a 2-3-fold increase in total lung reporter gene expression, along with improved distribution throughout the lung and increased activity in distal lung tissue ^17,18^. Importantly, PFCs do not alter the surface tension properties of the pulmonary surfactant because of their inert nature, thereby providing a stable environment for gas exchange ^19^. PFCs can also enhance lung retention by mitigating mucociliary clearance and limiting the rapid washout often seen with conventional aerosolized formulations due to the particles being inert ^14,20–22^. This prolonged retention time provides an extended window for cellular uptake and therapeutic activity, which is particularly advantageous for RNA interference (RNAi) therapeutics where sufficient intracellular accumulation is required for sustained RNA interference effects. Furthermore, the high oxygen solubility of PFCs can enhance various cancer treatments, including gene therapy, by improving tumor oxygenation and overcoming hypoxia-associated resistance ^23^. Certain PFC formulations can also potentially modulate immune responses to reduce the inflammatory response upon particle administration to the lungs ^24^. For instance, PFC administration decreased the release of leukotriene B4 and interleukin 6 in the bronchoalveolar lavage fluid of surfactant-depleted newborn pigs, playing a protective role against acute pulmonary inflammation ^25^. This potential modulatory effect of PFCs on acute inflammatory responses may be determined by the extent of PFC partitioning into the lipid bilayers of cellular membranes ^26^. Also, studies demonstrate that PFCs can block tumor necrosis factor-α-induced interleukin 8 release from alveolar epithelial cells and preventing neutrophil-mediated injury. This protection is achieved by PFCs acting as a mechanical barrier that prevents immune cells from penetrating their thin layers ^27–29^.

While PFCs hold great promise for pulmonary delivery applications, they present challenges due to the difficulty in formulating thermodynamically stable formulations suitable for clinical use ^30^. PFCs spread well on hydrophobic surfaces, however, they present formulation challenges due to the extremely high interfacial tension between PFCs and aqueous media, making it difficult to create stable PFC-water nanointerfaces without specialized surfactants. To address this, we proposed an inhalable RNA nanocapsule delivery platform utilizing PFCs surface-coated with an in-house developed amphiphilic polymer surfactant for enhanced stability and tumor targeting. Our design leverages PFCs’ unique properties to improve RNA delivery to tumors while maintaining stability during aerosolization and post-deposition. This PFC-based delivery system was engineered to facilitate the delivery of therapeutic miRNA that could reprogram the tumor microenvironment, thereby potentially increasing sensitivity to immune checkpoint blockade therapies through targeted modulation of immune responses in the tumor. In this study, we comprehensively characterized the nanocapsules for aerosolized delivery, modeled the impact of excipients on particle stability, evaluated their differential cellular uptake across various pulmonary cell types as well as pulmonary metastatic cells, assessed their safety profile, biodistribution, tumor retention, immune cell mediated clearance, and therapeutic efficacy through aerosolized delivery of therapeutic miRNA in mouse models of lung metastatic breast cancer. We also conducted a survival analysis and detailed immunomodulation studies to investigate the potential of our platform to enhance anti-tumor immune response within the tumor microenvironment and address existing limitations of conventional chemotherapeutic treatments (Figure 1).

**Figure 1.**
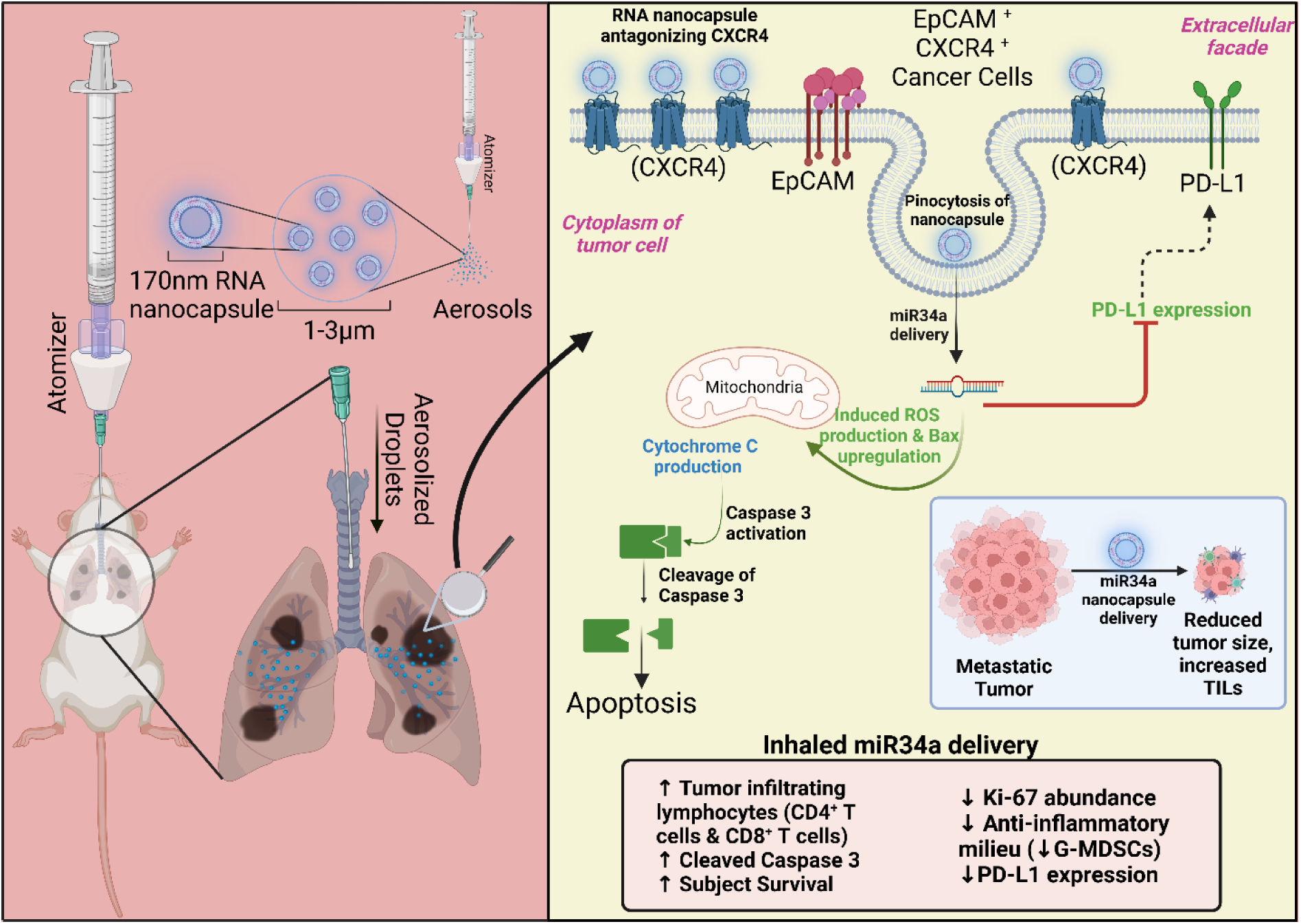
Schematic illustration of the mechanism of action of PFC-based RNA nanocapsules in the treatment of lung metastases. RNA nanocapsules encapsulating miR34a are formulated by mixing fixed ratios of nanoemulsion to miRNA. RNA nanocapsules are administered by microsprayer aerosolizer, intratracheally. CXCR4 is overexpressed on malignant lung metastatic cells which provides passive homing of the RNA nanocapsules to the sites of metastasis, aiding in the uptake and retention of RNA nanocapsules. Following endocytosis, the miR34a released from RNA nanocapsules suppresses PD-L1 expression and enhances cleavage of Caspase 3, thereby inducing apoptosis and alleviating metastatic tumors. miR34a nanocapsules exhibit multiple therapeutically beneficial mechanisms: we show that they induce apoptosis through increased levels of cleaved Caspase 3 and reduced Ki-67 abundance. Additionally, miR34a nanocapsules reduce the immunosuppressive milieu by reducing PD-L1 expression, allowing for increased tumor lymphocyte infiltration and reduction in granulocytic myeloid derived suppressor cells. *Created with Biorender.com*.

## Results and Discussion

Lung metastases affect nearly one-third of all cancer patients and pose a significant clinical challenge, particularly in malignancies like sarcomas, colorectal, lung, breast, and renal cancer ^31^. In breast cancer, lung metastases occur in about 27% of metastatic cases ^32^ and drastically impact disease outcomes, with overall mean survival of around 32 months following pulmonary metastasectomy ^33^. Lung metastases are often aggressive, exhibit high recurrence rates, and are influenced by hypoxic tumor microenvironment, which can promote cancer stem cell-like properties associated with chemoresistance and increased metastatic potential. Chemotherapy and pulmonary metastasectomy remain treatment options for select patients with limited lung metastases, offering potential for local control and improved survival ^34^. However, these options offer limited curative potential, as negative hormone receptor status, incomplete resection, early recurrence, and the presence of multiple metastases markedly worsen outcomes and patient prognosis ^34^. Thus, there is a critical need to develop novel approaches to treat lung metastasis to improve survival outcomes. Our previous work established poly(amidoamine) derived from a CXCR4 antagonist AMD3100 (PAMD) as an effective dual-function carrier to simultaneously inhibit CXCR4/SDF-1 signaling as well as achieve enhanced delivery of therapeutic nucleic acids ^35–37^. In this project, we report the formulation of oil-in-water PFOB nanoemulsions using the amphiphilic derivative of PAMD ^38,39^ as an emulsifier ^40,41^ and demonstrate its use as an inhalable therapy that outperforms conventional chemotherapies.

### Formulation and characterization of PFC RNA nanocapsules

PAMD and PAMD-C (MW 16.9 kDa) were synthesized and characterized as before (Supplementary Table 1, Supplementary Figure 1, Supplementary Figure 2, Supplementary Scheme 1). PAMD-C@PFOB oil-in-water nanoemulsions were prepared using a one-step ultrasonication method ^42^ using 1% v/v PFOB and 2 mg/mL PAMD-C.

The RNA nanocapsules were formulated by mixing PAMD-C@PFOB nanoemulsion and miRNA at different nanoemulsion-to-miRNA weight ratios (w/w) (Figure 2a). Colloidally stable nanocapsules with completely bound miRNA were formed at PAMD-C polymer to miRNA w/w ≥ 2. The hydrodynamic diameter (D_H_) of the nanocapsules decreased relative to the neat nanoemulsion without the miRNA, with values as low as 158 nm, which is ∼10-13% less than the DH of the neat nanoemulsion (Figure 2b). Reduction in nanoparticle size with increasing surface density of RNA likely stems from stronger electrostatic interactions with positive charged polymers, which has also been observed in other condensates ^43^. Additionally, while the zeta potential of the nanocapsules formulated at w/w of 2 and 4 (18 mV) was reduced slightly with respect to the neat nanoemulsion (21 mV), the residual positive zeta potential indicated that PAMD-C cationic moieties are present on the nanocapsule surface and not fully engaged in electrostatic interation with miRNA (Figure 2c). Hence, subsequent in vitro and in vivo studies were conducted using a w/w ratio of 4, so that the slight excess of CXCR4-binding moieties could enable targeting of CXCR4-overexpressing cells. CXCR4 binding capabilities of the RNA nanocapsules were confirmed in Supplementary Figure 3.

**Figure 2.**
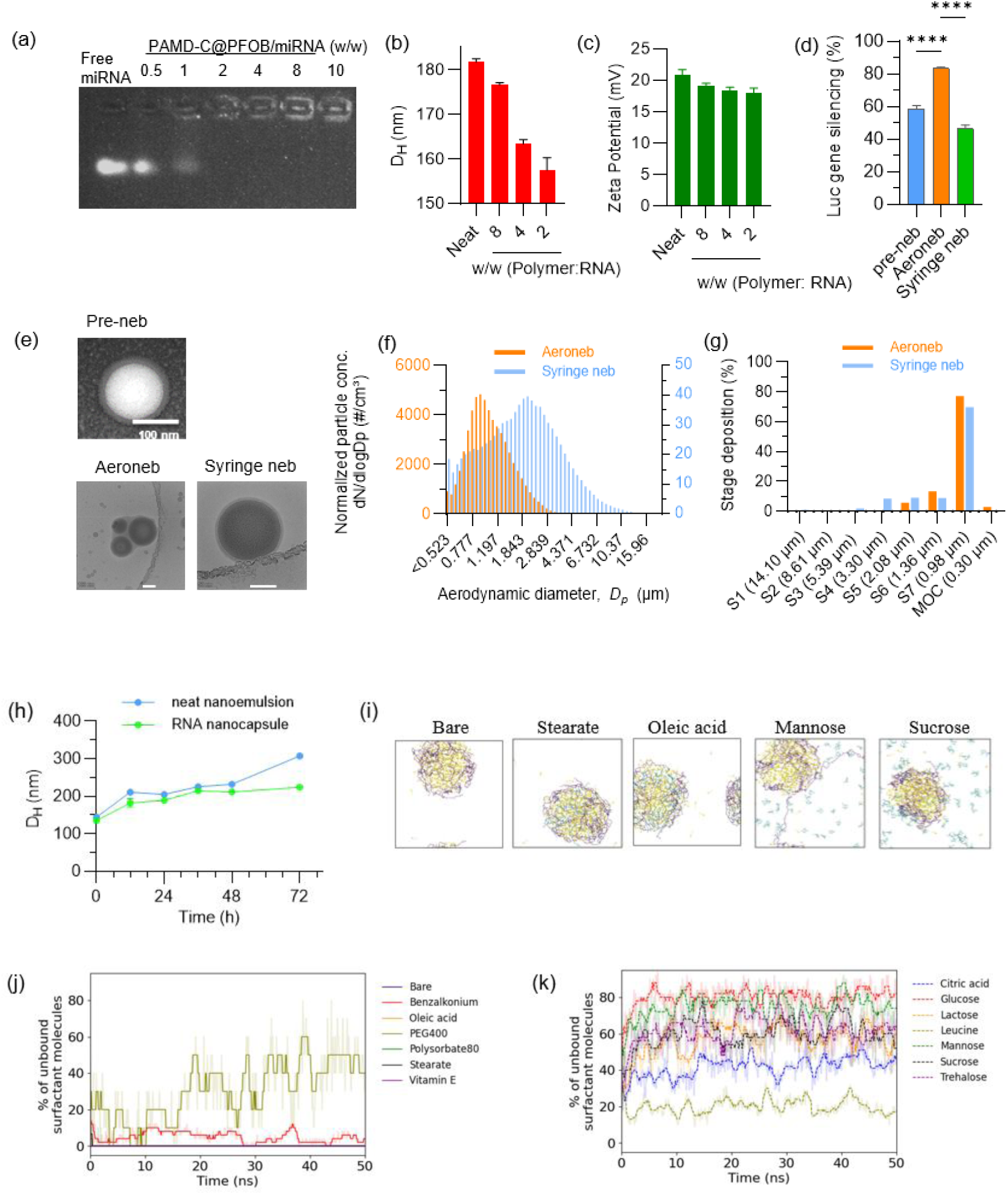
Formulation and characterization of PAMD-C@PFOB/miRNA nanocapsules. a) miRNA condensing ability of PAMD-C@PFOB nanoemulsion by agarose gel retardation assay. b) Mean hydrodynamic particle size and c) mean zeta potential of the RNA nanocapsules at different w/w ratios by DLS. d) In vitro luciferase gene silencing evaluation of PAMD-C@PFOB/siLuc nanocapsules in 4T1.Luc cells pre- and post-aerosolization. e) Representative TEM image of the RNA nanocapsule before aerosolization and representative CryoTEM images of the aerosolized RNA nanocapsules (w/w 4), scale bar = 100 nm in all three images. f) Aerodynamic particle size distribution from APS and (g) Fluorescence intensity from Cy3-labelled aerosolized RNA nanocapsules across different stage cut-off diameters of the NGI at 15 L/min. h) Colloidal stability of the nanoemulsions and the RNA nanocapsules at room temperature for three days. i) Snapshots from AAMD simumlations for various systems (PAMD-C + 1-fluorobutane + excipient). The first snapshot titled ‘Bare‘ denotes the reference system with PAMD-C and 1-fluorobutane. The other snapshots are titled using the name of the excipient included in the corresponding simulation. (j) and (k) show the evolution of the % of excipient molecules bound to the core 1-fluorobutane + PAMD-C particle. In (j-k), the solid lines are obtained after median filtering and the shaded regions denote the raw output data. The mean values and SD bars in (b-d) are obtained using n = 3 replicates each. **** p < 0.0001, *** p < 0.001, ** p < 0.01.

The nanoemulsions were aerosolized using two methods: (i) a vibrating mesh nebulizer (AeronebPro), denoted as Aeroneb and (ii) and a syringe microsprayer aerosolizer, denoted here as Syringe neb. Comparative physicochemical characteristics of the neat and aerosolized nanoemulsions are shown in Supplementary Figure 4. Luciferase gene knockdown study was done in 4T1.Luc cells to study preservation of the RNA functionality post nebulization. While Luc gene silencing observed in Syringe neb and pre-neb samples were comparable, Aeroneb showed a ∼ 41% increase in Luc gene silencing activity (Figure 2d). This improved transfection efficiency observed with Aeroneb aerosolized nanocapsules likely stems from their reduced zeta potential (Supplementary Figure 4 d), which represents an optimal balance of positive charge. While positive charge is necessary for interaction with negatively charged cell membranes, excessively high positive charge can cause cytotoxicity. The moderately positive zeta potential of Aeroneb-delivered nanocapsules appears to strike an ideal balance that maintains sufficient electrostatic attraction for cellular uptake, thereby enhancing transfection efficiency ^44^. The RNA nanocapsules exhibited a spherical core-shell architecture as visualized by TEM, where the PFOB core was stabilized by a PAMD-C/miRNA shell layer (Figure 2e). This structural organization allowed for the presentation of PAMD-C and its CXCR4-binding moieties on the emulsion surface, thereby maintaining the positive surface charge required for miRNA binding. Cryo-TEM revealed distinct morphological changes in the nanoparticle structure following nebulization (Figure 2e). Both pre- and post-nebulized samples exhibited discrete, well-defined nanoparticles with clear core-shell boundaries. RNA nanocapsules were safe up to 150 nM RNA when tested in vitro using cell viability assay (Supplementary Figure 5 a, b). In vitro uptake of the RNA nanocapsules in 4T1.Luc cells with strong cytoplasmic colocalization of PAMD-C@PFOB-Cy3 and miRNA-FITC was validated in Supplementary Figure 6.

In order to effectively deliver the nanocapsules to lung tumors, it is essential that the aerodynamic diameter of the aerosolized nanocapsules is appropriate for targeting the specific lung region where the tumor is located. For tumors in the peripheral lung regions where many lung adenocarcinomas develop, particles of 1-5 μm diameter are required to achieve optimal deposition. The droplet size distribution obtained from APS is shown in Figure 2f. The droplets of RNA nanocapsule solution aerosolized using Aeroneb are dominated by a single log-normal mode. The total distribution has a peak aerodynamic diameter (Dp) value of ∼1 µm and a tail that extends out to 3 µm. The RNA nanocapsules aerosolized using the syringe neb are composed of multiple log-normal modes. The total distribution has a peak Dp value of ∼2.1 µm and a tail that extends out to 12 µm. In NGI analysis, both the Aeroneb and Syringe neb samples have a clear deposition spike at stage S7 (0.98 µm), and reduced depositions at stages S4, S5, and S6 ( Figure 2g). Overall, the aerodynamic size of the RNA nanocapsule using both nebulizers is conducive to deep lung deposition. Results from in vivo expeirments in further sections confirm these hypotheses.

The size of the RNA PFC nanocapsules increases only by 20% over three days at room temperature, which is negligible compared to the rapid 113% increase in D_H_ values for the PAMD-C@PFOB nanoemulsion (Figure 2h). To study the impact of excipient molecules on particle agglomeration and thereby its shelf-life, AAMD simulations were performed for various systems. Since the actual particle sizes of ∼170 nm are beyond the scope of AAMD simulations, the impact of excipients was modeled as a self-assembly problem. Figure 2i shows the simulation snapshots for few representative systems, where each system consists of PAMD-C, 1-fluorobutane (approximation for PFOB), and an excipient. The ‘bare’ system without any excipient self-assembles to form a nearly spherical particle with a ∼2.5 nm radius. Most 1-fluorobutane molecules (>99.5%, yellow) assemble to form the core and all PAMD-C molecules (purple) are deposited on the core. Upon adding stearate/oleic acid, the excipient molecules (green) are bound to the core particle. In contrast, mannose/sucrose molecules are randomly dispersed in the simulation cell and show very weak interactions with the core particle. Further, clustering analysis presented in Figure 2(j,k) shows the qualitative differences between the two sets of excipients. Figure 2j shows that the % unbound excipient molecules upon adding oleic acid, polysorbate 80, stearate, and vitamin E is 0 (same as the bare system which does not have any excipients), confirming that all excipient molecules are bound to the particle. Other excipients in Figure 2k have large % unbound values. Overall, our results suggest that weak excipients such as mannose, sucrose, etc. may not inhibit particle agglomeration. However, strong excipients like stearate and oleic acid may prevent or delay the agglomeration of neighboring particles, as supported by previous simulations for solid hydride particles^45,46^. Other results from clustering analyses and details regarding the simulation setup are shown in Supplementary Figure 7 and Supplementary Table 4 respectively.

### In vivo lung uptake and retention of aerosolized RNA nanocapsules

To evaluate the intrapulmonary distribution and retention of the RNA nanocapsules in the lungs and lung tumors, we conducted studies in both healthy and lung metastatic breast cancer (LMBC) mice. We administered dual-labeled RNA nanocapsules (Cy3-PAMD-C@PFOB/Cy5.5 miRNA) at a dose of 20 µg miRNA and 80 µg PAMD-C in 60 µL. The dose was selected based on previous report ^47^. The dual fluorescent labeling enabled tracking of both the delivery vehicle and the therapeutic cargo. Enhanced accumulation of Cy5.5-labeled miRNA in the lungs was observed compared to other major organs, for both healthy and LMBC mice (left and right panels in Figure 3a). At 24 h, the average radiance in the LMBC mice lungs is 1.5 times higher than the average radiance in the healthy mice lungs, signaling longer retention of the nanocapsules by the tumor mice. Moreover, in healthy mice, the significantly decreased intensity of the Cy5.5 signal in the lungs at 48 h indicates that the miRNA is cleared from the lungs after 24 h (Figure 3b). On the other hand, there is no significant decrease in the Cy5.5 signal intensity in the LMBC mice lungs between 24 and 48 h (Figure 3b). This enhanced retention in tumor-bearing lungs likely results from sequestration by the mononuclear phagocyte system (MPS) of the lung, which has been shown to retain PFC nanoparticles ^48^.

**Figure 3.**
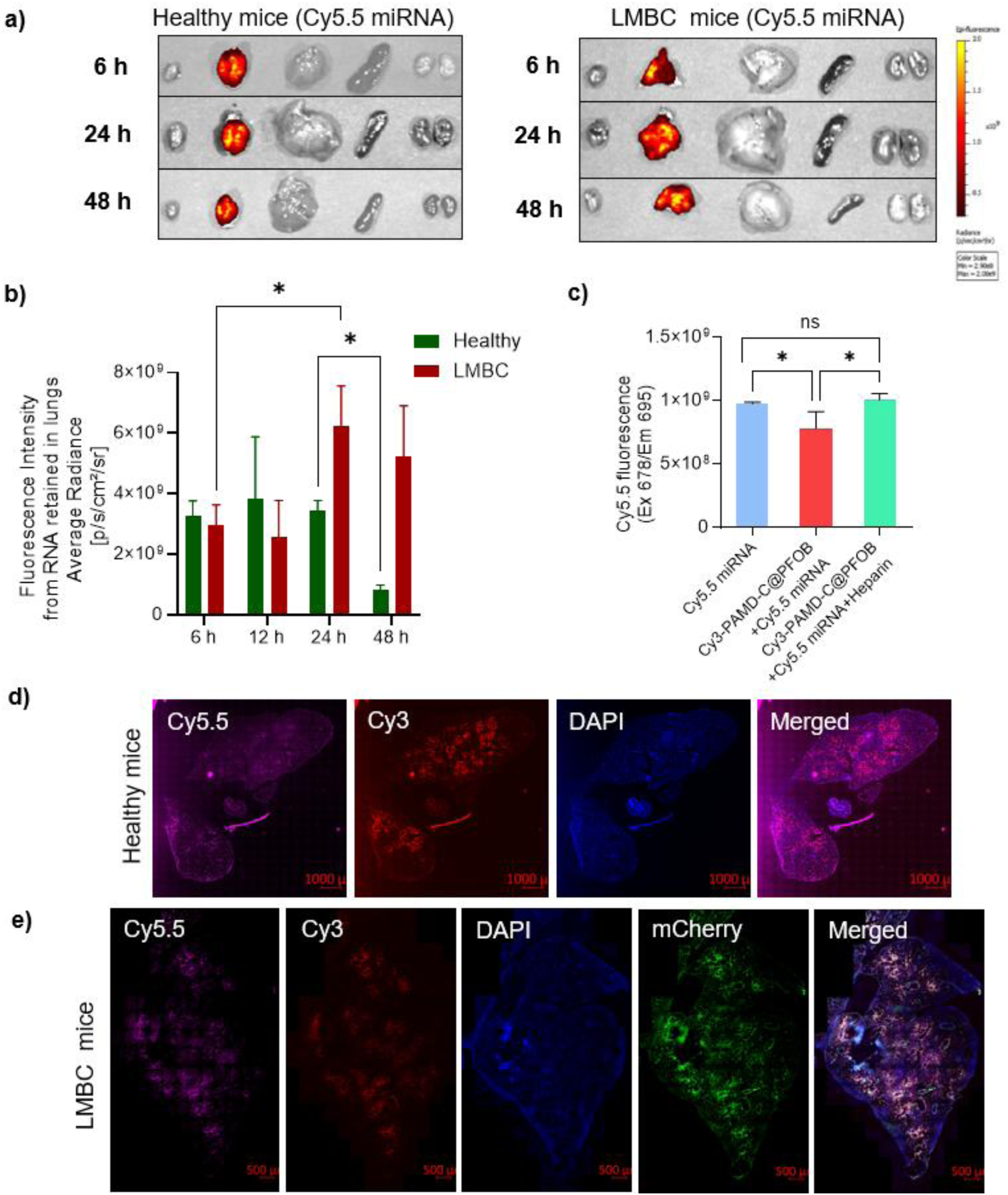
Biodistribution of PAMD-Ch@PFOB/miRNA nanocapsules in healthy and LMBC mice. (a) Representative ex-vivo fluorescence images of major organs at different time points post aerosolized administration of nanocapsules in healthy and LMBC mice. b) Mean fluorescence intensity of the miRNA retained in the lungs of healthy and LMBC mice at different time points post-aerosolized administration of the nanocapsules (c) Fluorescence intensity from Cy5.5 miRNA with and without complexation to Cy3-PAMD-C@PFOB nanoemulsion. (d) Representative confocal images of lung sections 24 h post administration of labeled RNA nanocapsules in healthy mice. Scale bar = 1000 μm . (e) Representative confocal images of lung sections 24 h post administration of labeled RNA nanocapsules in LMBC mice injected with 4T1-Luc-mCherry cells. Scale bar = 500 μm. n = 3 per group . * p < 0.05, ns-not significant.

Administering miRNA to the lungs should result in the highest levels of fluorescence immediately following administration, followed by gradual reduction due to clearance from the lungs. However, the data in Figure 3b shows comparable or even higher fluorescence intensities at 24 h and 48 h, compared to the values at 6 h and 12 h. This trend points towards a controlled release of the miRNA payload from the nanocapsule due to their disassembly in the cells. To test this hypothesis, we examined the nanocapsules under three conditions: i) Cy5.5 miRNA, ii) nanocapsule with Cy5.5 miRNA, and iii) nanocapsule with Cy5.5 miRNA and Heparin. As seen in Figure 3c, the fluorescence signal is significantly quenched when the miRNA is in the nanocapsule, relative to bare miRNA (blue vs red bars), likely due to the close proximity of the Cy 3 and Cy 5.5 fluorophores. Upon adding heparin to the nanocapsules, the fluorescence signal increases to levels comparable to that of the bare miRNA, demonstrating heparin-mediated release of miRNA from the nanocapsule. This observation demonstrates that the miRNA is not instantaneously released from the carrier nanocapsule after administration, as evident from the relatively weak average radiance at 6 h and 12 h. At longer times (24 h and 48 h), the miRNA is released from the nanocapsule once taken up by the lung cells, especially in tumor mice. Detailed results from flow cytometry experiments in the following sections show miRNA uptake in various lung cell populations.

To assess lung penetration, we analyzed the intra-pulmonary distribution of the labeled RNA nanocapsules 24 h post aerosolized administration, using confocal microscopy. We observed co-localization of both fluorescent markers in epithelial cells surrounding the alveoli in healthy mice (Figure 3d). This colocalization suggests that the miRNA maintained their structural integrity throughout the delivery process while undergoing gradual release at the target site.

To specifically evaluate delivery of the RNA nanocapsules to the tumors in the lung, we utilized 4T1.mCherry cells to generate the lung metastatic tumors. Confocal microscopy revealed clear colocalization between the RNA nanocapsules and mCherry-expressing tumor cells, confirming successful delivery into the metastatic lesions (Figure 3e).

### Inflammatory response and cellular distribution of the RNA nanocapsules in lung tissue of healthy and LMBC mice

To evaluate inflammatory response to the RNA nanocapsules in healthy mice, a 40 cytokine array was performed 12 h post-administration (Supplementary Figure 8). Figure 4a shows that only two cytokines, CXCL1 and TIMP-1, were highly upregulated compared to placebo when threshold for fold change was set at 1.5. The upregulation of TIMP-1 may be linked to the interaction between TIMP-1 and the CXCR4 signaling axis. TIMP-1 has been shown to influence CXCR4-mediated pathways, which play a role in immune cell recruitment, activation and tissue remodeling ^49^. Disrupting this axis by the CXCR4 antagonizing nanocapsules can enhance their expression and affect immune cell homeostasis in the lungs. Additionally, CXCL1 has been implicated in promoting neutrophil migration and activation, processes that are often transient during inflammatory responses. CXCR4 antagonists, such as AMD3100, have been shown to modulate chemokine levels, including CXCL1, in a time-dependent manner ^50^. This suggests that the upregulation of TIMP-1 and CXCL1 likely represents a transient immune activation phase triggered by the CXCR4 antagonistic properties of our delivery system, potentially serving as a protective mechanism to regulate inflammation and promote tissue repair in healthy mice.

**Figure 4.**
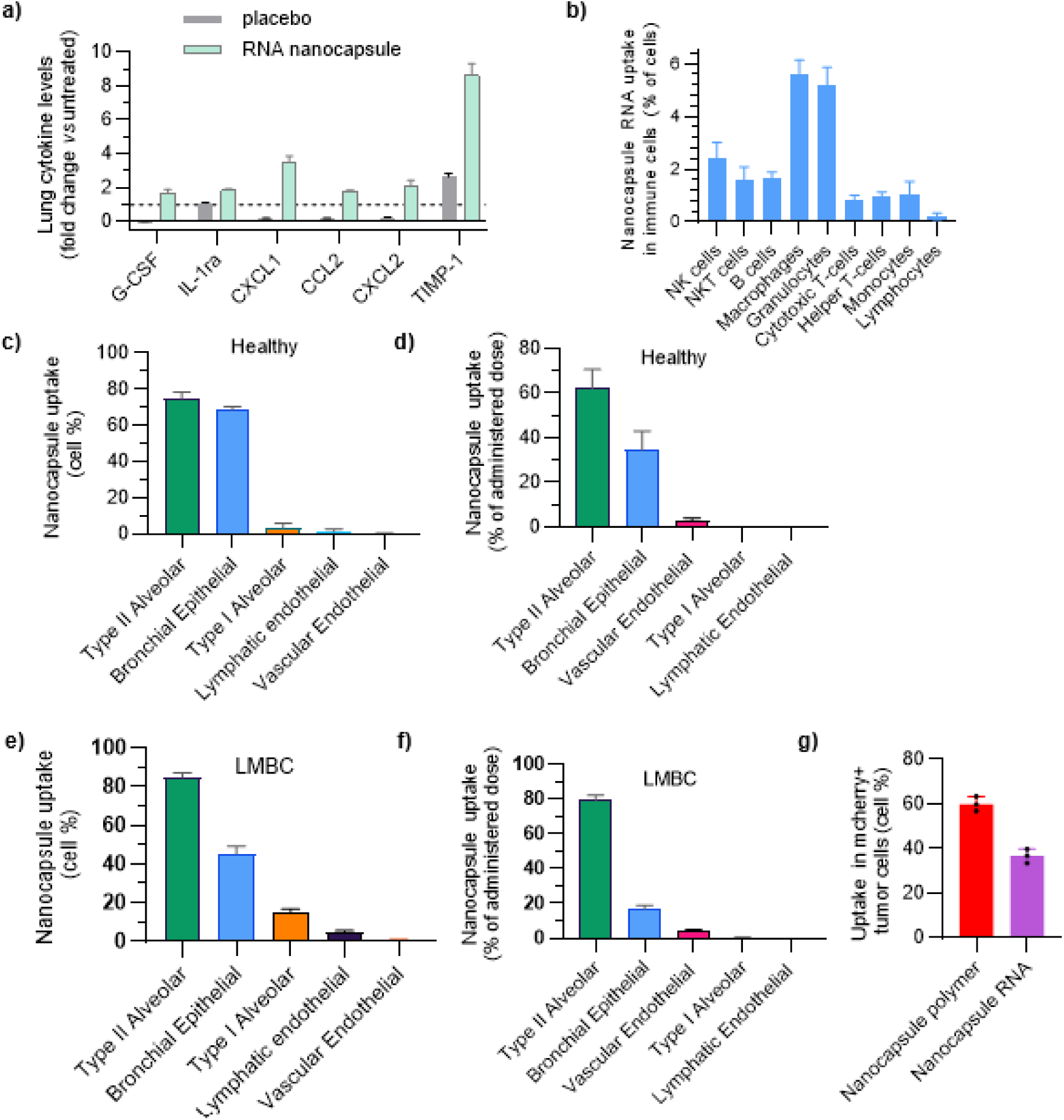
Cellular distribution and safety assessment of miRNA nanocapsules in healthy and LMBC-mCherry mouse lungs. a) Pulmonary immune response profile following RNA nanocapsule administration. Analysis of lung tissue protein lysates from healthy controls, placebo-treated (DI H_2_O 60 µl), and RNA nanocapsule-treated (20 µg miRNA) mice at 12h post-administration (n = 3). A panel of 40 immune mediators was evaluated, including cytokines (TNF, IL-6, IL-12p70, IL-1β, IFNγ, IL-2, IL-17, IL-5, IL-16, TIMP1, TREM1, IL-23, IL-10, IL-1ra, IL-13 and IL-4), chemokines (CCL1, CCL2, CCL3, CCL4, CCL5, CCL11, CCL12, CCL17, CXCL1, CXCL2, CXCL9, CXCL10, CXCL11, CXCL12 and CXCL13), growth factors (G-CSF, GM-CSF, M-CSF, IL-3 and IL-7) and cell adhesion molecules (CD54 and complement component 5a) (Data are presented as mean fold change relative to healthy controls. Raw images are shown in Supplementary Figure 8. (b) Flow cytometric analysis of immune cell populations showing uptake of Cy5.5-labeled nanocapsule miRNA in healthy mouse lungs 4 h post-administration. Distribution of Cy5.5 labeled nanocapsule miRNA in healthy mouse lung CD45-populations, showing (c) percentage uptake per cell type and (d) percentage of total administered dose. (e,f) Similar analysis in LMBC mouse lungs showing (e) cellular uptake and (f) total dose distribution in CD45-subtypes. (g) Uptake of Cy3-labeled nanocapsule polymer and Cy5.5-labeled nanocapsule miRNA by 4T1-mCherry tumor cells isolated from the lungs of LMBC mice.

The rapid clearance of therapeutic formulations by immune cells poses a significant challenge for effective pulmonary delivery. Phagocytic cells such as macrophages and other lung resident immune cells rapidly recognize and engulf foreign materials, reducing the bioavailability and efficacy of treatments intended for non-immune target cells. To evaluate our delivery system’s ability to evade this clearance mechanism, we first conducted flow cytometric analysis to evaluate uptake of Cy5.5-labeled miRNA by different CD45+ pulmonary immune cells in healthy mouse lungs. Results revealed that only 2-6 % of the administered dose was taken up by various pulmonary immune cells indicating effective evasion of immune recognition (Figure 4 b). This limited uptake by pulmonary immune cells is advantageous for applications targeting non-immune cells that necessitate prolonged localization in the intended target cells to achieve optimal therapeutic efficacy.

The distribution of miRNA across lung epithelial and endothelial cells (CD45-) was also analyzed. We found that ∼80% of type II alveolar cells and 70% of bronchial epithelial cells have taken up the nanocapsules, while other analyzed cell types demonstrated less than 5% uptake (Figure 4c). Analysis of the total miRNA dose distribution revealed that type II alveolar cells accounted for approximately 60% of the administered dose, while bronchial epithelial cells which line the conducting airways of the lungs from trachea to the terminal bronchioles contributing 30% (Figure 4d). The remaining cell types, including type I alveolar, vascular and lymphatic endothelial cells, collectively accounted for less than 10% of total uptake. This preferential accumulation in type II alveolar and bronchial epithelial cells can be attributed to their robust endocytic activity ^51^. Conversely, the limited uptake observed in other cell types likely stems from differences in their glycocalyx composition, affecting nanoparticle binding and internalization ^52^. Similarly, in LMBC mice, most of the miRNA was taken up by type II alveolar cells (Figure 4e and f), with relatively low uptake by other cell-types. Lastly, Figure 4g shows that the polymer and miRNA uptake by 4T1.mcherry cells of LMBC mice was 60% and 38%, respectively, demonstrating RNA nanocapsule delivery to the tumor cells.

### Anti-tumor effect of aerosolized miR-34a PFC nanocapsules

In light of the resistance of metastatic breast cancers to immune checkpoint blockade therapy, we investigated the therapeutic potential of miR-34a nanocapsules using a lung metastasis model. miR-34a, which is downregulated in various cancers including lung and breast cancer, acts as a tumor suppressor and regulates expression of immune checkpoint proteins like PD-L1 ^53^. We hypothesized miR-34a could effectively treat lung metastases through its pleiotropic effects on multiple cellular pathways, including PDL1-mediated immune regulation, cell survival, metastasis, and tumor progression networks ^53,54^. Following tumor implantation, the animals were treated with aerosolized RNA nanocapsules on days 6, 10, and 14 (Figure 5a). Controlled animals received placebo and nab-paclitaxel (nab-ptx) chemotherapy (10 mg/kg) via intravenous administration on days 5 and 9.

**Figure 5.**
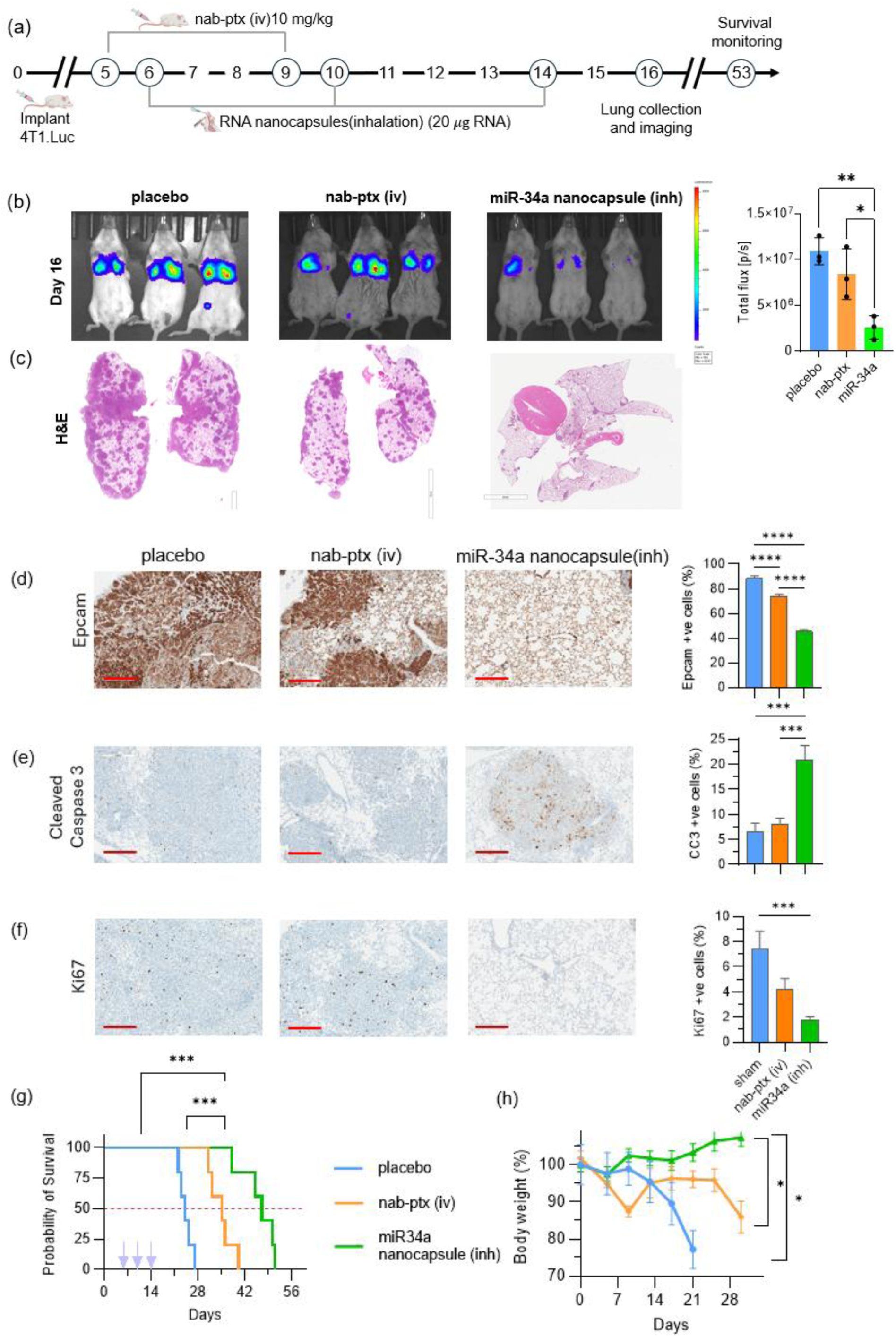
In vivo antitumor efficacy of the miR-34a nanocapsules in LMBC mice. (a) Experimental timeline showing the dosing regimen for LMBC mice (b) In vivo bioluminescence imaging and quantification of total bioluminescent flux from the lungs of LMBC mice post different treatments. (c) H&E staining of lung tissues collected on day 16 (scale bar, 5mm). Representative images of immunohistochemical staining across different treatment groups and Percentage of positively stained area for (d) Epcam (e) Cleaved Caspase 3 (CC3) (f) Ki67. n = 3 mice per group. Areas in the sections having metastatic tumors only were quantified. ImageJ was used for quantification of the images. Three fields for each section was quantified. Scale bar = 300 µm. (g) Survival curves after different treatments (n=5) analyzed using log-rank test. Median survival: placebo (24 days), nab-ptx (35 days), miR34a nanocapsule (47 days) (h) Plots representing percentage body weight change in the different treatment groups (n = 5). **** p < 0.0001, *** p < 0.001, ** p < 0.01, * p < 0.05

In vivo bioluminescence imaging on day 16 revealed differential treatment responses across groups. While the nab-ptx group showed moderate tumor inhibition, the miR-34a group demonstrated significant tumor inhibition with nearly five fold reduction in total bioluminescent flux compared to untreated mice (Figure 5b). H&E staining of harvested lungs further supported the observation, showing drastically reduced metastatic burden in the miR-34a group , with smaller and fewer lung metastases compared to nab-ptx group (Figure 5c). Immunohistochemical analysis further validated the therapeutic efficacy, revealing that miR-34a treatment induced a substantially larger reduction in Epcam+ epithelial-derived carcinoma marker cells compared to nab-ptx (78% vs 40%), indicating enhanced tumor regression and reduced epithelial characteristics (Figure 5d). Conversely, cleaved caspase 3 positive (CC3+) cells, indicative of apoptosis were significantly increased in both treatment groups, with miR-34a demonstrating a stronger pro-apoptotic effect compared to nab-ptx (Figure 5e). Ki67, a marker for cell proliferation, was significantly reduced in both nab-ptx and miR-34a groups compared to placebo, with miR-34a group demonstrating only 2% Ki67+ cells — a two-fold and four-fold decrease compared to nab-ptx and placebo group respectively (Figure 5f). In addition to the therapeutic outcomes resulting from miR34a treatment, we hypothesize that the CXCR4 antagonism by the nanocapsules may have contributed to reduced tumor burden ^55^.

Survival analysis further demonstrated the superior therapeutic efficacy of miR-34a nanocapsules. While nab-ptx treatment modestly extended median survival from 24 to 35 days, miR-34a nanocapsule therapy significantly doubled the survival of the placebo group to 47 days, despite cessation of treatment after day 14 (Figure 5g). This substantial survival benefit, achieved with just three doses of the nanocapsule, suggests that additional treatment cycles beyond day 14 could potentially yield further improvements in survival outcomes.

Body weight trends also showed major differences between the various treatment groups. Placebo group experienced a steep decline in body weight by ∼25% within three weeks. Similarly, the body weight of the nab-paclitaxel treated mice was reduced by 15% within ten days – likely due to the effects of chemotherapy given on days 5 and 9, and later after three weeks, due to recurrent tumors at later stages. In contrast, the mice receiving miR-34a nanocapsules showed steady or gradually increasing body weights over four weeks (Figure 5h). This initial progress was observed after just three doses, with body weight monitored for only 30 days. As no additional doses were administered, any residual tumor cells in the miR34a group could have potentially proliferated, leading to death at around day 50.

### Treatment effect on immunological effectors in the tumors and anti-tumor immunity

To investigate the impact of miR-34a nanocapsule therapy on immune modulation within the TME, we analyzed the composition of a few immunological effectors in metastatic lungs. We hypothesized that the combined effect of miR-34a treatment and PAMD-C mediated CXCR4 blockade would synergistically promote T cell infiltration while reducing immunosuppressive cells. IHC analysis revealed that miR-34a treatment significantly decreased CD45+ cells by ∼50 % compared to nab-ptx and placebo groups (Figure 6a). This is reflective of the marked decrease in granulocytic myeloid-derived suppressor cells (G-MDSCs) within the lungs (Figure 6e). Since G-MDSCs represent a major component of the CD45+ infiltrate that increases proportionally with tumor burden in the 4T1 model , this reduction suggests an alleviation of the tumor burden ^56^.

**Figure 6.**
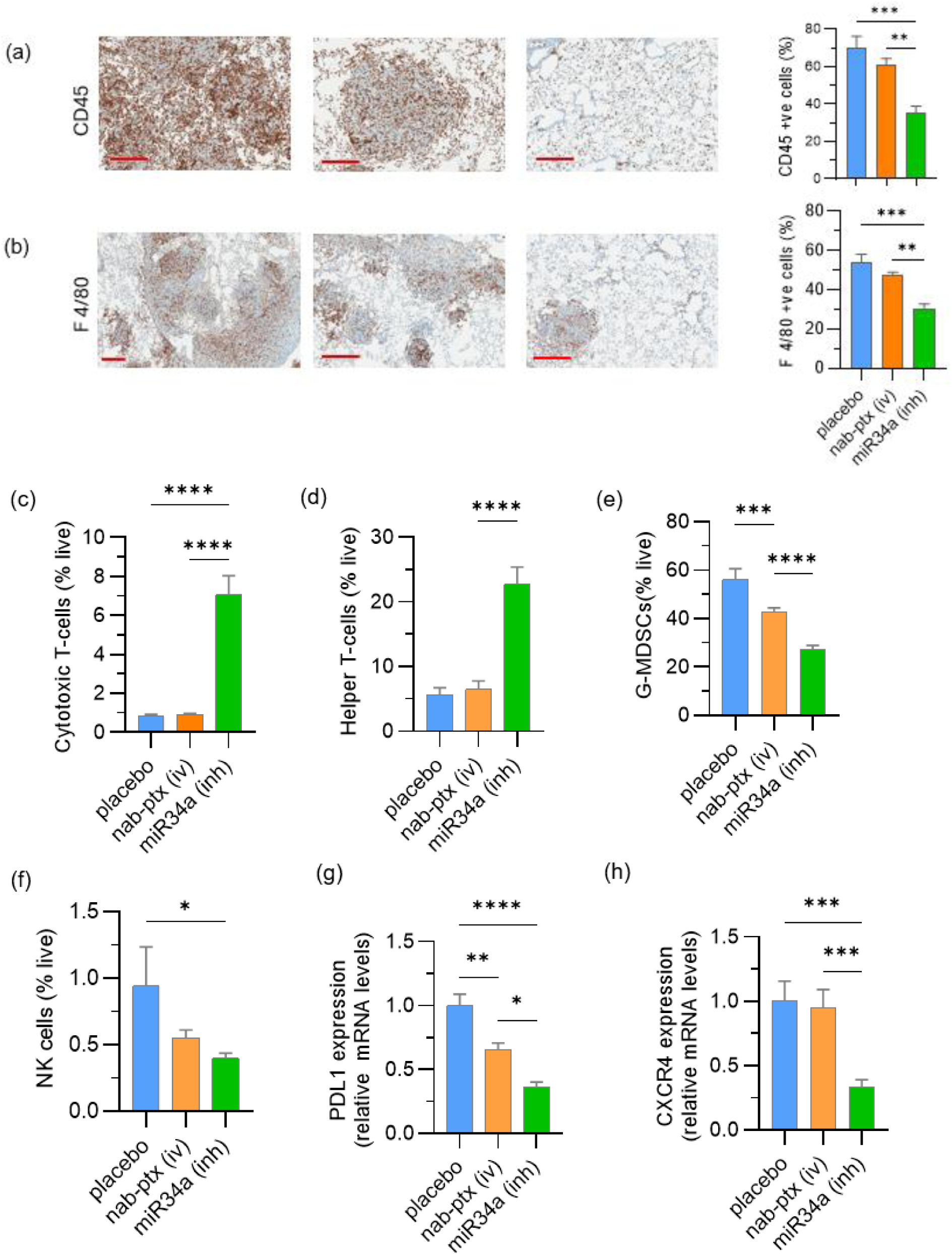
Effect of miR34a nanocapsule treatment on the levels of select immunological effectors in LMBC mice. Representative IHC staining and Percentage of positively stained area for (a) CD45 and (b) F4/80. Levels of (c) Cytotoxic T-cells, (d) Helper T-cells, (e) G-MDSCs, and (f) NK cells across different treatment groups. Pulmonary mRNA levels of (g) PDL1 and (h) CXCR4. GAPDH was used as an internal control. n = 3, **** p < 0.0001, *** p < 0.001, ** p < 0.01, * p < 0.05

F4/80+ tissue-resident immune cells (typically of monocytic lineage such as macrophages and monocytes) were reduced by ∼10% in the nab-ptx group and ∼ 50% in the miR34a group, compared to placebo (Figure 6b). Since F4/80+ cells promote the induction of inducible regulatory T cells (iTregs) ^57^ and anti-inflammatory cytokine production that fosters oncogenesis, this substantial decrease in the miR-34a group likely indicates a shift towards a more immune-permissive environment, conducive to immune-mediated tumor clearance.

Flow cytometry revealed a significant increase in CD8+ T cell infiltration in miR-34a group, increasing from ∼1% (of total live cells) in placebo and nab-ptx groups to nearly 8% (Figure 6c). This increase is indicative of the environment becoming more permissive to immune-mediated tumor clearance, consistent with the expected effects of CXCR4 axis antagonism enhancing CD8+ T cell infiltration into tumors by disrupting the CXCL12/CXCR4 chemokine gradient (that normally restricts T cell trafficking) and alleviating immunosuppression ^58^. Similarly, CD4+ helper T cells increased substantially from ∼5% in control groups to ∼22% in the miR-34a group (Figure 6d), suggesting a reinvigorated immune response given their crucial role in supporting CD8+ cytotoxic T cell function within the TME ^59^.

Further, the miR34a group demonstrated significant reduction in granulocytic myeloid-derived suppressor cells (G-MDSCs) from ∼ 60% in placebo and 45% in nab-ptx to 30% (Figure 6e). Since G-MDSCs utilize immunosuppressive mechanisms to inhibit anti-tumor immune responses ^60^, this reduction suggests a broad attenuation of immunosuppressive signals ^61^, contributing to increased T cell infiltration and activation ^62,63^. Natural killer (NK) cells were also significantly reduced (>50%) in the miR-34a group compared to placebo (Figure 6f), likely due to miR-34a-mediated downregulation of ULBP2, an NK cell immune-receptor ligand essential for tumor recognition and lysis ^64^. This reduction stems from miR-34a’s inherent properties, independent of the delivery system’s efficacy.

Critically, these observed immune changes correlated with decreased PD-L1 mRNA levels in the miR-34a group (Figure 6g). 4T1 tumors produce granulocyte-colony stimulating factor (G-CSF), a pro-granulopoietic growth factor that drives G-MDSC proliferation^65^, which in turn fosters PD-L1 upregulation contributing to immune suppression and leukocyte exhaustion. The significant reduction of PD-L1 suggests decreased tumor burden and alleviation of the immunosuppressive TME. This reduction mirrors the observed increases in T cell infiltration (Figure 6c, d) by removing a major inhibitory signal that would otherwise limit anti-tumor activity.

Also, miR-34a nanocapsule treatment decreased CXCR4 mRNA levels (Figure 6h), which reduces terminally exhausted T cells (T_ex_) while promoting functional CD8+ T cell infiltration^66^ and directly inhibiting CXCR4-driven cancer progression and metastasis. However, CXCR4 downregulation may potentially create a feedback mechanism wherein subsequent nanocapsule doses experience diminished targeting efficiency due to reduced receptor availability, highlighting the importance of optimized dosing schedules and potential benefits of combination targeting strategies for sustained therapeutic efficacy. Together, these findings suggest that inhalable miR-34a nanocapsules achieve therapeutic efficacy through their dual action: enhancing anti-tumor immunity while disrupting CXCR4-dependent processes that support tumor growth and metastases.

In summary, we developed a safe and non-invasive inhalation platform that addresses the challenge of delivering RNA therapeutics effectively to lung tumors by navigating mucus barriers and evading immune clearance. Unlike previous inhalable delivery systems that show poor tumor accumulation and rapid elimination, our platform uniquely combines the biocompatibility of perfluorocarbons with active CXCR4-mediated tumor targeting, achieving unprecedented tumor uptake (60% of tumor cells) and extended retention (>48 hours). The therapeutic efficacy of these nanocapsules encapsulating miR34a is demonstrated through potent pulmonary downregulation of PDL1 and CXCR4, markedly improved reduction in lung metastatic tumor burden, enhanced localized anti-tumor immunity, and nearly doubled lifespan compared to standard chemotherapy after just three doses. Moreover, the nanocapsules demonstrate potent passive targeting of type II alveolar and bronchial epithelial cells, thereby opening doors for the treatment of other lung diseases of epithelial origin. Collectively, these results open new avenues for the use of PFCs as nucleic acid carriers in inhaled genetic medicines with superior performance and clinical translation potential.

## Methods

### Materials

All cell culture materials were obtained from Thermo Fisher Scientific, including Roswell Park Memorial Institute-1640 (RPMI-1640, catalog #11875093), Phosphate Buffered Saline (PBS, catalog #10010023), Fetal Bovine Serum (FBS), trypsin, penicillin, and streptomycin. Chemical reagents were sourced as follows: hexamethylenebisacrylamide (HMBA) from Pfaltz & Bauer (Waterbury, CT); 1-bromoheptadecafluorooctane (perfluorooctyl bromide , PFOB), N,N-diisopropylethylamine (DIPEA), heparin sodium salt (CAS #9041-08-1, catalog #H3149), and Cholesteryl chloroformate from Sigma-Aldrich (St. Louis, MO); and AMD3100 from A2B Chem LLC (catalog #AD67785). Nucleic acid reagents were acquired from various suppliers: negative control miRNA (miScr), carboxyfluorescein (FAM)- and Cy5.5-labeled miRNA, and microRNA 34a (mmu-miR-34a-5p) from Horizon Discovery. Cy3-NHS was procured from Lumiprobe Corporation (catalog #21020). The Proteome Profiler Mouse Cytokine Array Kit, Panel A (catalog #ARY006) was purchased from R&D Systems. Reverse-transcription PCR primers were obtained from Sigma-Aldrich and Sino-Biological Inc. Antibodies for immunophenotyping were sourced from BD Biosciences, BioLegend, and ThermoFisher Scientific.

### Synthesis of PAMD-Cholesterol@PFOB nanoemulsions

Polymeric AMD3100 (PAMD) was synthesized via Michael-type addition using equimolar concentrations (1.9 mmol) of AMD3100 and HMBA as previously described ^38^. The reactants were dissolved in methanol/water (7/3 v/v) and reacted at 37°C under nitrogen for 4 days in darkness. Subsequently, an additional one-tenth of the initial AMD3100 quantity was added and reacted for another day to consume any unreacted acrylamide. The reaction mixture was purified by dialysis (3.5 kDa MWCO) against methanol for 2 days, followed by dialysis against deionized water for an additional day. The final product was obtained by lyophilization at 38% yield. PAMD structure was confirmed by ¹H-NMR (Supplementary Figure 1), and molecular weight was determined by gel permeation chromatography (GPC) using an Agilent 1260 Infinity LC system with a TSKgel G2000PWXL-CP column in 0.1 M sodium acetate buffer (pH 5).

PAMD-Cholesterol (PAMD-C) was synthesized with 17% cholesterol substitution(Supplementary Table 1). PAMD (310 mg) was dissolved in anhydrous methylene chloride (10 mL) with DIPEA (242.58 mg), and cholesteryl chloroformate (68.18 mg) in anhydrous methylene chloride was added dropwise over 1 hour on ice. The reaction proceeded for an additional 24 hours under stirring. The product was obtained by solvent evaporation and washed three times with diethyl ether to remove unreacted cholesteryl chloroformate. The product was then dissolved in ethanol/water (1:1 v/v), and the pH was adjusted to 4.0 using 1.25 M HCl. Following dialysis (3.5 kDa MWCO) against ethanol/water (1:1 v/v) for 2 days and deionized(DI) water for 1 day, the product was lyophilized (85% yield). PAMD-C structure was confirmed using ¹H-NMR (Supplementary Figure 2). The intensity of integration of the cholesterol methyl group to the aromatic phenylene protons of PAMD was used for determining mol% and weight% of cholesterol in PAMD-C (Supplementary Figure 2, Supplementary Table 1). The molecular weight of PAMD-C was determined using GPC (Supplementary Table 1).

Fluorescently labeled polymers were synthesized by reacting PAMD-C with Cy3-NHS ester (1% w/w) for 24 h at 40°C in darkness, followed by dialysis against distilled water for one day and lyophilization. PAMD-C@PFOB nanoemulsions were formulated by adding 20 μL PFOB to 2 mL PAMD-C (2 mg/mL in DI water) and ultrasonicating using a probe-type ultrasonic processor (Hielscher, UP200ST) equipped with a 2 mm diameter probe, with energy restricted to 8000 W at 80% output amplitude.

### RNA nanocapsule preparation and characterization

RNA nanocapsules were formulated by mixing PAMD-C@PFOB nanoemulsions with 0.5 µg miRNA at different weight-to-weight (w/w) ratios of the PAMD-C polymer and miRNA in 10 mM HEPES buffer (pH 7.4) followed by incubation at room temperature for 30 min. The binding of miRNA to PAMD-C@PFOB was analyzed by a gel retardation assay. Nanocapsules prepared at different PAMD-C@PFOB to miRNA w/w ratios were loaded into the wells of a 2% agarose gel containing a SYBR^TM^ Safe dye, which binds to unencapsulated miRNA. Gel electrophoresis was performed for 15 min at 100 mV in 0.5x Tris/borate/EDTA buffer. miRNA encapsulation was visualized using an E-Gel^TM^ Imager (Invitrogen). The zeta potential and hydrodynamic diameter of the RNA nanocapsules were determined by dynamic light scattering (DLS) using the Zetasizer (Malvern Pananalytical). Nanocapsule morphology was examined by Transmission electron microscopy (TEM) using Nanovan® (Nanoprobes, catalog #2011) as a negative stain as well as by Cryo-TEM.

### Aerosolization

Two aerosol generation systems were tested — microsprayer aerosolizer (PenWu Device for Mouse, BioJane, Cat# BJ-PW-M) and Aeroneb® Lab Nebulizer Small VMD (Kent Scientific Corporation, Cat# NEB-1100). For aerosol characterization, the RNA nanocapsules were prepared with 20µg miRNA and 80µg PAMD-C polymer( at w/w 4 from 2 mg/mL stock) and diluted in DI water in a ratio of 1:5 for aerosolization purposes as dilution led to lesser aggregation of the particles during aerosolization. For animal studies, the RNA nanocapsules were administered undiluted through the microsprayer.

### Characterization of aerosolized RNA nanocapsules

To probe the aerodynamic size, the nanocapsules were aerosolized in an upwelling flow chamber, with an Aerodynamic Particle Sizer (APS) and a Next Generation Impactor (NGI) connected to the sample port. The upwelling vacuum flow rate was set at 50 L/min, and the flow rate leaving the side sampling port flow rate was set at 16 L/min. The APS sampled at 1 L/min while the NGI sampled at 15 L/min ( Supplementary Figure 9 a). The vacuum pulled for a total of 20 min, including a background sample for 10 min and a sample collection following aerosolization for another 10 min. The samples were aerosolized up to 1 minute for AeroNeb lab nebulizer or burst of aerosols from microsprayer aerosolizer and recorded for 10 minutes. APS (model 3321, TSI, Shoreview, MN, USA) was used to assess aerosol size and concentration. APS data was recorded on a connected laptop that used the Aerosol Instrument Management (v.10.3) software (TSI Inc., Shoreview, MN, USA).

NGI (MSP, Shoreview, Minn, USA) was used to determine the size of the collected particles. The NGI is a seven stage cascade impactor consisting of 8 collection cups with the following size range cut-offs: 14.10 µm, 8.61 µm, 5.39 µm, 3.30 µm, 2.08 µm, 1.36 µm, 0.98 µm, and a final micro-orifice collector (MOC) of 0.3 µm (Supplementary Figure 9 b). Following each experimental run, impacted particles in the collection cups were rinsed with 10 mL of deionized water, and 200 µL was collected in a Qubit assay tube for fluorometry testing.

### Aerosol Size and Quantification

To determine impacted particle size and quantity, Cy3 labelled PAMD-C@PFOB nanoemulsion was used. Following the sampling timeframe, fluorescent quantification was done with the Qubit 4 Fluorometer (Q33226, Invitrogen by Thermo Fisher Scientific, Wilmington, DE, USA). The fluorometer’s green emission spectra (510 -580 nm) were used based on the excitation peak of Cy3∼ 554nm to calculate the relative fluorescence units (RFU). Emission readings were entered into Excel for analysis. The R^2^ values and slope of the curve were used to determine the fluorescein concentration in mg/mL. The RFU values were fit to the curve using the following equation: 𝑦 = 30030𝑥 + 2609.2 ; R^2^ = 0.9606

### Cell culture

4T1-Luc [Ubigene] (RRID: CVCL_C8UY) and NCI-H1975-Luc [Ubigene] (RRID: CVCL_C8W1) cell lines were obtained on January 2021 and March 2022 respectively. Cells were cultured in RPMI-1640 media with 10% FBS and 1% penicillin-streptomycin. 4T1-Luc-mCherry cells (RRID: CVCL_C8UZ) were a kind gift from Dr. Shyamala Maheswaran from Massachusetts General Hospital Harvard University in June, 2024 and they were cultured in Gibco RPMI-1640 (A10491-01) (ATCC modification 30-2001) with 10% FBS and 1% penicillin-streptomycin. Recombinant human osteosarcoma epithelial cells U2OS expressing EGFP-tagged CXCR4 receptor was obtained in October, 2018 from Thermo Fisher, USA (#K1779, RRID:CVCL_ZK12) and were cultured in DMEM high glucose medium supplemented with 10% FBS, 2 mM L-Glutamine, 1% Penicillin-Streptomycin and 0.5 mg/mL G418. Cells were cultured in a humidified incubator at 37 °C with 5% CO_2_.

### In vitro cytotoxicity

In vitro cytotoxicity of nanoemulsions and nanocapsules was assessed by CellTitre-Blue (CTB) cell viability assay (Promega, catalog # G8080). 4T1-Luc and H1975-Luc cells were seeded in 96-well plates at a density of 10,000 cells/well and incubated overnight. The next day, cells were treated with increasing concentrations of the nanoemulsion and nanocapsules (formulated with increasing concentrations of miRNA) and incubated for 24 h. Then, cell viability was evaluated by CTB assay following the manufacturer’s protocol. The half maximal inhibitory concentration (IC_50_) values were calculated from a dose-response analysis in GraphPad Prism.

### Luciferase gene silencing by aerosolized RNA nanocapsules in vitro

4T1.Luc cells were seeded in a 48-well plate at a density of 3 × 10⁴ cells per well and incubated overnight. RNA nanocapsules (PAMD-C@PFOB/siLuc) containing siLuc( *Silencer*™ Firefly Luciferase (GL2 + GL3) siRNA , Thermo Fisher, Cat# AM4629) or negative control siRNA at w/w 4, 150 nM siRNA per well, were prepared and subjected to aerosolization using the Aeroneb lab nebulizer and the microsprayer aerosolizer. Both pre- and post-aerosolized RNA nanocapsules were then used to transfect the cells in serum-free medium for 6 hours. Following transfection, the medium was replaced with fresh medium containing 10% FBS and kept for 48 hours. Subsequently, cells were washed with PBS, lysed using cell lysis buffer, and luciferase activity was quantified using the Promega Luciferase Assay System (Cat. No. E1500).

### CXCR4 inhibition assay

CXCR4 inhibitory activity by PAMD-C@PFOB was assessed by CXCR4-redistribution assay in U2OS cells stably expressing EGFP-bound CXCR4 receptor as reported in previous studies^67^. U2OS cells were seeded in black clear-bottomed 96 well plates at a density of 15,000 cells/well and incubated overnight. Next day, the cells were washed twice with 100 μL of assay buffer and incubated with increasing concentrations of PAMD-C@PFOB nanoemulsion for 30 mins. Next, 10 nM of stromal cell-derived factor 1 (SDF-1, Millipore Sigma Catalog # GF344) was added to the cells followed by incubation for 1 h. The cells were fixed with 10% formalin for 20 mins followed by three washes with PBS and then visualized using a fluorescence microscope (EVOS® FL). CXCR4 receptor internalization was measured via a Cellomics ArrayScan V high-content analysis plate reader, which detected fluorescent spots formed when CXCR4-EGFP fusion proteins moved into endosomes. Here, SDF-1 functioned as the agonist, while the CXCR4-specific inhibitor AMD3100 was used as a reference antagonist.

### Intracellular localization and quantification of cellular uptake of the nanocapsules

To assess subcellular localization of the RNA nanocapsules via confocal microscopy, 4T1-Luc cells were plated on 15 mm glass-bottomed cell culture dishes overnight at a density of 10,000 cells/well. Next day, cells were transfected with nanocapsules prepared using Cy3-labelled PAMD-C@PFOB and FITC-labelled miRNA (w/w 4, 150 nM miRNA) for 6 h. Cells were washed thrice with PBS, and the nuclei were stained with 4′,6-diamidino-2-phenylindole (DAPI). Finally, cells were fixed using 10% formalin before imaging on a confocal microscope (Zeiss LSM 800 with Airyscan). For quantification of uptake of the labelled RNA nanocapsules using flow cytometry, 4T1-Luc cells were seeded in 6-well plates overnight and the above-mentioned treatments were given the next day for 6 h. Cells were then washed thrice with PBS, detached with TrypLE (Gibco), and analyzed by flow cytometry on a FACSCalibur instrument (BD Biosciences). Free FITC labelled miRNA was used as a control.

### Animal models of lung metastatic breast cancer (LMBC)

All animal experiments were conducted using 8-week-old female Balb/c mice obtained from Charles River Laboratories. All experiments were performed according to an approved protocol by the Institutional Animal Care and Use Committee(IACUC) having protocol no 19-018-03FC of the University of Nebraska Medical Center. The LMBC model was established by tail vein injection of 2x10^5^ 4T1-Luc-mCherry cells in 100 µl sterile PBS per mouse. This modified cell line was used to track metastasis and tumor progression through both bioluminescent imaging (via the luciferase protein expression) and fluorescent visualization (via mCherry protein expression). LMBC model using the 4T1 cells (a highly metastatic mammary cancer cell line) serves as a model for stage IV human breast cancer. Studies show that mice implanted with 4T1 cells developed progressively growing tumors that spontaneously metastasized to the lungs, ultimately resulting in death from metastatic disease in all animals ^68^. The lung metastasis was confirmed at different days post implantation starting at day 5, by IVIS bioluminescence imaging after intraperitoneal injection of Luciferin substrate (3 mg per mouse). The mice were provided with a standard diet and had unrestricted access to water.

### Biodistribution of RNA nanocapsules

Biodistribution was assessed in healthy and LMBC mice model at different time points post administration of nanocapsules. On day 12, post tumor implantation, lung metastasis was confirmed by *in vivo* bioluminescence imaging. Then the microsprayer aerosolizer was used to administer aerosolized RNA nanocapules (60 μL) formulated with Cy3-labelled PAMD-C@PFOB and Cy5.5-labelled miRNA(Dharmacon) (20 μg miRNA per mouse, w/w=4) in mice (n=3 per group). Mice were euthanized at different times (6 h, 12 h, 24 h and 48 h) post administration of the nanocapsules and major organs were collected for *ex vivo* imaging on a Xenogen IVIS 200 system.

### Intra-pulmonary distribution of labelled RNA nanocapsules in healthy and LMBC mice

On day 12, lung metastasis was confirmed in LMBC mice by *in vivo* imaging on a Xenogen IVIS 200 system detecting fluorescence from mcherry cells. Then mice in the healthy as well as LMBC group received an equivalent volume of the labelled nanocapsules as mentioned above. To assess intra-pulmonary distribution of the nanocapsules, lungs were perfused and embedded in Tissue-Tek® optimal cutting temperature (O.C.T.) Compound (Sakura® Finetek Catalog # 4583), sectioned, stained with DAPI, and then fluorescence from Cy3, Cy5.5 and mcherry was visualized on a Zeiss Axioscan 7 microscope.

### In vivo therapeutic efficacy

The therapeutic efficacy of RNA nanocapsules was evaluated in LMBC model. Mice were randomly assigned to three treatment groups (n=15 per group): (1) Placebo receiving saline (i.v.) and DI water (aerosol), (2) NAB Paclitaxel (i.v.), (3) aerosolized PAMD-C@PFOB/miR34a. NAB-Paclitaxel was administered intravenously on days 5 and 9 at a dose of 10 mg/kg body weight in 100 μL sterile PBS, totaling two doses. RNA nanocapsule treatments were administered intratracheally using a microsprayer aerosolizer (PenWu, BioJane) on days 6, 10, and 14, totaling 3 doses. Each nanocapsule dose consisted of 20 μg miR34a formulated with PAMD-C@PFOB nanoemulsion (w/w 4) and 10% 10 mM HEPES in a total volume of 60 μL. Body weights were recorded throughout the study. On day 16, three mice per group were injected intraperitoneally with 3 mg D-luciferin in 100 μL PBS for bioluminescence imaging to assess tumor burden. Five mice from each group were monitored for survival until day 53 without any additional treatment. On day 16, multiple analyses were performed: four mice per group were sacrificed for flow cytometric analysis of the tumor immune microenvironment using lung single-cell suspensions; three mice per group were used for RNA extraction from lung tissue to confirm relative mRNA levels of PDL1, a target of miR34a , via RT-PCR; and the remaining three mice per group had their lungs perfused with formalin fixed in 4% paraformaldehyde for 24 hours, embedded in paraffin, and sectioned for H&E and IHC staining.

### Pulmonary single cell isolation and flow cytometry analysis of nanocapsule uptake

We employed flow cytometry to quantify the uptake of Cy5.5-labelled miRNA in mouse lung epithelial, endothelial, and hematopoietic lineage cells. Additionally, this method also allowed us to assess immunomodulation within the lung tissue following various treatments.

### Epithelial & endothelial cell (CD45-) isolation

At the time of lung tissue harvest, the pulmonary circulation was flushed by injecting 10 mL of cold 1x PBS containing 500 U/mL heparin(150 μL from 10,000 u/ml) through the left ventricle of the heart. The lung was inflated with 1.5 mL cold Dispase solution (Corning, Catalog # 354235) by instilling a catheter through the trachea. This was followed by perfusion with 200 µL of 1% low melting point (LMP) agarose (CAS# 9012-36-6, GoldBio). The lungs were immediately placed in ice and incubated for 2 min, followed by incubation for 45 minutes in 5 mL of Hank’s Balanced Salt Solution without Calcium and Magnesium (HBSS) (Thermo Fisher, Catalog # 14175095) at room temperature. The lungs were minced using scissors and transferred to a gentleMACS™ C tube (Miltenyi Biotec Catalog # 130-093-237) containing 1 mL of Dulbecco’s Modified Eagle Medium (DMEM) and 100 μL of 1 mg/mL DNase I (Sigma-Aldrich, Catalog # 11284932001).

### CD45+ cell isolation

Collagenase Type IV (Sigma, CAS #9001-12-1) was prepared at a concentration of 700 mg per 500 mL of Dulbecco’s Modified Eagle Medium (DMEM). Following the pulmonary circulation flushing procedure described previously, the lungs were minced using scissors and transferred to a gentleMACS™ C tube. Each tube contained 1 mL of the prepared DMEM with Collagenase IV, 100 μL of DNase I (1 mg/mL), and 30 μL of Heparin (from 10,000 U/mL).

The C tubes were placed upside down in the gentleMACS™ Dissociator (Miltenyi Biotec, Catalog # 130-093-235) and the tissue was dissociated for 8 seconds to form a single cell suspension, which was immediately transferred to ice. An additional 4 mL of DMEM was added, and the tubes were kept in a rotating incubator at 150 rpm for 30 minutes at 37°C. Then added 5 mL of 1x PBS containing 4 mM EDTA at pH 7.4 (Teknova Catalog # P0203) and the cells were sequentially filtered through pre-separation filters of 70 μm (Miltenyi Biotec, Catalog # 130-095-823) and 40 μm (Corning Cat # 352340). The cell suspension was centrifuged at 1500 rpm for 10 minutes. After decanting the supernatant, 1 mL of ACK Lysis Buffer (Quality Biological, Catalog # 118-156-721) was added to the cells and vortexed gently to lyse the RBCs. This mixture was kept for 2 minutes, after which 2 mL of DMEM was added. It was again vortexed gently, and the cells were centrifuged at 1500 rpm for 5 minutes. The supernatant was decanted, and the cells were resuspended in 500 μL of FACS Buffer (1X PBS + 2% FBS + 0.1% NaN_3_). The cells were counted in the TC10™ automated cell counter and the antibodies used for the flow cytometry staining are shown in Supplementary Table 2. The gating strategy used to separate the pulmonary immune and epithelial/endothelial cells as well for immunophenotyping in the therapeutic study are shown in Supplementary Figure 10 and Supplementary Figure 11, respectively.

### Cytokine Array

To assess inflammation in lung tissue post administration of the RNA nanocapsules, a cytokine array was conducted in 8-week-old female Balb/c healthy mice. The study included three groups: one receiving aerosolized PAMD-C@PFOB/miRNA nanocapsules, a healthy control group, and a placebo control group receiving aerosolized DI water. Twelve hours after administering the formulations, lungs were harvested, protein lysates prepared by homogenizing the lung tissue in RIPA lysis buffer containing protease and phosphatase inhibitor cocktails (Thermo Fisher Scientific). Protein concentration was determined using the Bradford reagent. Approximately 300 μg of pooled protein extract from lung tissue was used for cytokine quantification. The antibody-based protein array (Proteome Profiler: Mouse Cytokine Array, R&D Systems, catalog number: ARY006) was employed according to the manufacturer’s instructions. Densitometry of cytokine spots on the membrane was performed using Image Lab 6.1 software (Bio-Rad) with the volumetric method to determine cytokine expression levels after background subtraction.

### Reverse transcription–polymerase chain reaction (RT-PCR)

Total RNA was extracted from lung tissues using TRIzol reagent (Invitrogen #15596026). The RNA was then converted to cDNA using a high-capacity reverse transcription kit from Applied Biosciences (#43-688-14). RT-PCR analysis was conducted on a Roto-Gene Q (Qiagen) thermocycler using a iTaq Universal SYBR Green Super mix (Bio-Rad #1725121) following the manufacturer’s protocol. Relative mRNA levels were calculated using the Ct values obtained. All primer sequences used are listed in Supplementary Table 3.

### Histological analysis

Mouse lung tissues from the different treatment groups were perfused and fixed in 4% paraformaldehyde and then stored in 75% ethanol. Then the lung tissues were embedded in paraffin, sectioned and stained with hematoxylin and eosin (H&E) using standard protocols. For immunohistochemistry (IHC), antigen retrieval was performed on the tissue sections by immersing them in heated citrate buffer (pH 6.0) for 20 min. 3% hydrogen peroxide was used to block endogenous peroxidase activity, followed by incubation in 5% normal goat serum to prevent non-specific binding. The tissue sections were then incubated at 4°C overnight with primary antibodies, followed by 1 h incubation at room temperature with HRP-conjugated secondary antibody. All slides were imaged using a Leica Aperio CS2 scanning system and images were analyzed using a Aperio Image Scope Software (Leica Biosystems).

### Molecular Dynamics (MD) Simulations

Interaction potentials for all molecules in this study were parametrized using the CHARMM code . PFOB was approximated using 1-fluorobutane since the CHARMM force field was not directly available for fully fluorinated alkanes. All-atom molecular dynamics (AAMD) simulations were run using the GROMACS code . Input configurations (9 nm × 9 nm × 9 nm) were generated using CHARMM-GUI ^69^. All systems were neutralized at a pH of 7 using Na+ and Cl-ions. AAMD simulations were run for 50 ns (2 fs timestep) in the NPT ensemble at 303.15 K and 1 bar. Snapshots were saved every 100 ps. Details for each system including number of molecules and their structure are shown in the Supplementary Table 4. Visualization and clustering analysis was done using OVITO ^70^.

### Statistical analysis

Results are presented as mean ± standard deviation. Statistical differences between three or more groups were analyzed using one-way ANOVA with Tukey’s post-hoc test, while Student’s t-test was employed for analyzing differences between two groups. Survival curves were analyzed using the log-rank test. A p-value < 0.05 was considered statistically significant, and all statistical analyses were conducted using GraphPad Prism 9.

## Data Availability Statement

The data that support the findings of this study are available from the corresponding author upon reasonable request.

## Acknowledgements

This research was supported by funds from the University of Nebraska Medical Center and grants P30 GM127200, R01 CA235863, R01 DK120533, and R01 DK124095. We gratefully acknowledge the University of Nebraska Medical Center’s Advanced Microscopy Core Facility, Flow Cytometry Core Facility, and Tissue Sciences Core Facility for their support. We would also like to thank Holly C. Britton, Dr. Craig L. Semerad, Amy J. Nelson, and Dr. Sudipta Panja, for their contributions to this work. The Core Facility receives partial support from NIGMS INBRE (P20 GM103427) and COBRE (P30 GM106397) grants, as well as support from NCI for The Fred & Pamela Buffett Cancer Center Support Grant (P30 CA036727). Major instrumentation has been provided by the Office of the Vice Chancellor for Research, The University of Nebraska Foundation, the Nebraska Banker’s Fund, and by the NIH-NCRR Shared Instrument Program. The content and interpretations presented in this publication are solely the responsibility of the authors.

## Author contributions

K.S. and D.O. conceived the idea, performed data analysis, and drafted the manuscript. K.S., A.M., A.S., B.R.P., N.K., L.D., C.J.W., M.A.K., C.M.J., and A.R.R. conducted the experiments. J.A.P., J.L.S., and J.E.T. supervised the experimental work. All authors reviewed, edited, and approved the final version of the manuscript.

## Additional Information

Supplementary Information

