## Supplementary Information for "Inhalable Perfluorocarbon RNA Nanocapsules Bypass Immune Clearance While Targeting Lung Epithelial and Lung Tumor Cells"

**Supplementary Table 1–** Characterization of PAMD-C copolymers.

| Polymer/  Copolymer | Mn (kg/mol) | Mw (kg/mol) | Polydispersity(M_w_/M_n_) | Cholesterol content (wt% in copolymer) |
| --- | --- | --- | --- | --- |
| PAMD | 12.3^b^ | 13.9^b^ | 1.1^b^ | - |
| PAMD-C | 12.2^b^ | 16.9^b^ | 1.3^b^ | 17^ab^ |

^a^From ^1^H-NMR

^b^From GPC

**Supplementary Table 2** **–** List of antibodies used for flow cytometry.

| **Panel Name** | **Antibody** | **Clone** | **Fluorochrome** | **Vendor** | **Catalog #** |
| --- | --- | --- | --- | --- | --- |
| Myeloid | CD11c | HL3 | BUV395 | BD Biosciences | 564080 |
| Myeloid | Ly6C | HK1.4 | BV785 | BioLegend | 128041 |
| Myeloid | CD11b | M1/70 | BV711 | BioLegend | 101242 |
| Myeloid | CD45 | 30-F11 | BV605 | BioLegend | 103155 |
| Myeloid | Ly6G | 1A8 | BV480 | BD Biosciences | 746448 |
| Myeloid | CD24 | M1/69 | RB705 | BD Biosciences | 757888 |
| Myeloid | MHCII (I-A/I-E) | M5/114.15.2 | AF488 | BioLegend | 107615 |
| Myeloid | F4/80 | QA17A29 | PE-Cy7 | BioLegend | 157307 |
|  | Fixed Viability | X | Ghost Dye Red 780 | CYTEK | 13-0865-T100 |
| Myeloid/LSK | CD49b | DX5 | APC | BioLegend | 108909 |
| Myeloid/LSK | CD3 | 145-2C11 | APC | BioLegend | 100311 |
| Myeloid/LSK | CD19 | 1D3/CD19 | APC | BioLegend | 152409 |
| Myeloid/LSK | Ter-119 | Ter-119 | APC | BioLegend | 116211 |
| Myeloid/LSK | Sca-1 | D7 | BV421 | BioLegend | 108128 |
| Myeloid/LSK | CD117 (cKit) | 2B8 | BUV615 | ThermoFisher | 366-1171-80 |
| T Cell | CD8a | 53-6.7 | BUV395 | BD Biosciences | 563786 |
| T Cell | CD25 | PC61.5 | BV786 | BioLegend | 102051 |
| T Cell | CD4 | GK1.5 | BV711 | BioLegend | 100447 |
| T Cell | CD19 | 1D3 | BV480 | BD Biosciences | 566167 |
| T Cell | CD3 | 145-2C11 | RB705 | BD Biosciences | 570648 |
| T Cell | DX5 (CD49b) | DX5 | PE-Cy7 | BioLegend | 108921 |
| Epi/Endothelial | CD326 (Epcam) | G8.8 | BUV395 | BD Biosciences | 740281 |
| Epi/Endothelial | CD31 | 390 | BV785 | BioLegend | 102435 |
| Epi/Endothelial | CD29 | HM Beta1-1 | BV711 | BD Biosciences | 740708 |
| Epi/Endothelial | Podoplanin | 8.1.1 | BV421 | BioLegend | 127423 |
| Epi/Endothelial | CD24 | M1/69 | RB705 | BD Biosciences | 757888 |
| Epi/Endothelial | MHCII (I-A/I-E) | M5/114.15.2 | AF488 | BioLegend | 107615 |

**Supplementary Table 3** **–** Primer Sequences

| **Primer Name** | **Company** | **Catalog #** | **Primer Sequence** | |
| --- | --- | --- | --- | --- |
|  |  |  | **Forward** | **Reverse** |
| GAPDH | Sino Biological | MP200537 | NA | NA |
| PD-L1/  B7-H1 | Sino Biological | MP200010 | NA | NA |

**Supplementary Table 4** **–** MD simulation setup

| **Excipient** | **#molecules** | **Excipient** | **#** | **Excipient** | **#** | **Excipient** | **#** |
| --- | --- | --- | --- | --- | --- | --- | --- |
| Benazlkonium | 50 | Leucine | 100 | Polysorbate 80 | 10 | Vitamin E | 10 |
| Citric acid | 100 | Mannose | 50 | Stearate | 15 |  |  |
| Glucose | 50 | Oleic acid | 50 | Sucrose | 50 |  |  |
| Lactose | 50 | PEG400 | 10 | Trehalose | 50 |  |  |

Each system contained 300 1-fluorobutane molecules and 10 PAMD-C molecules. The number of excipient molecules is shown below (larger count for smaller molecules and vice-versa). The empty space was filled with water molecules.

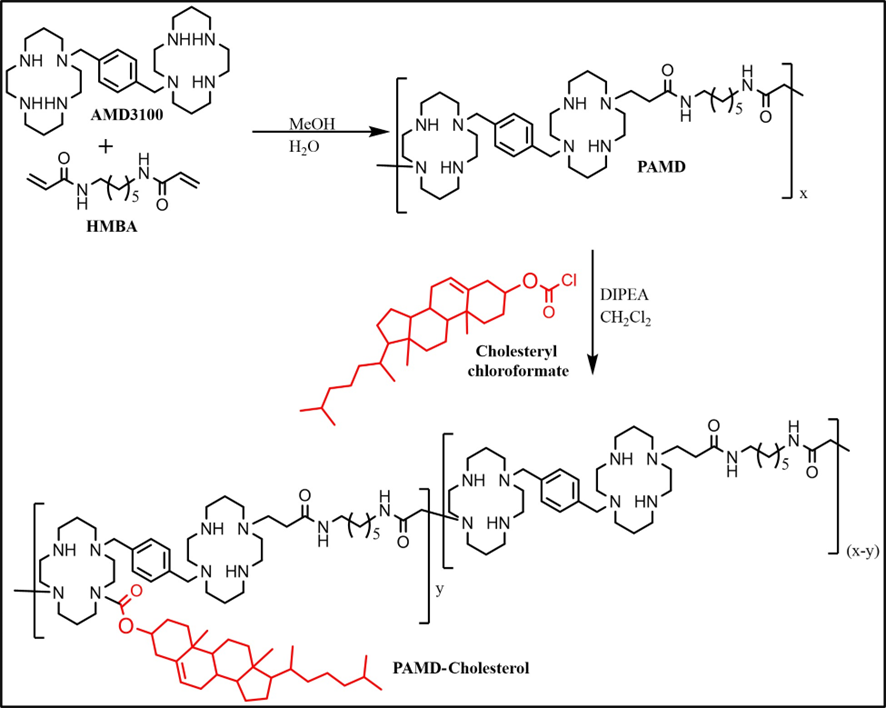

**Supplementary Scheme 1: Synthesis of PAMD-C.** Any of the secondary amine groups (-NH-) in the cyclam ring can participate in the Michael-type addition reaction.

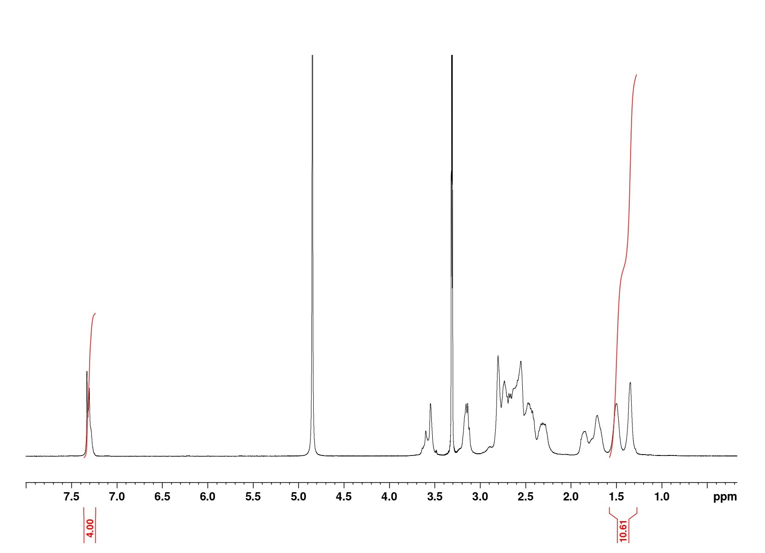
 **Supplementary Figure 1.** Typical ^1^H-NMR spectrum of PAMD in methanol D4

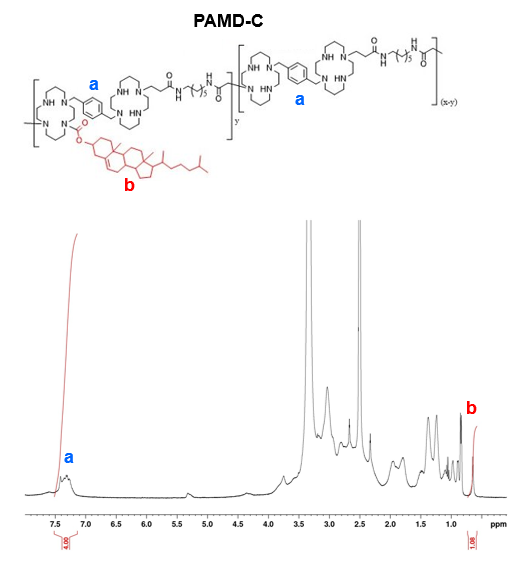

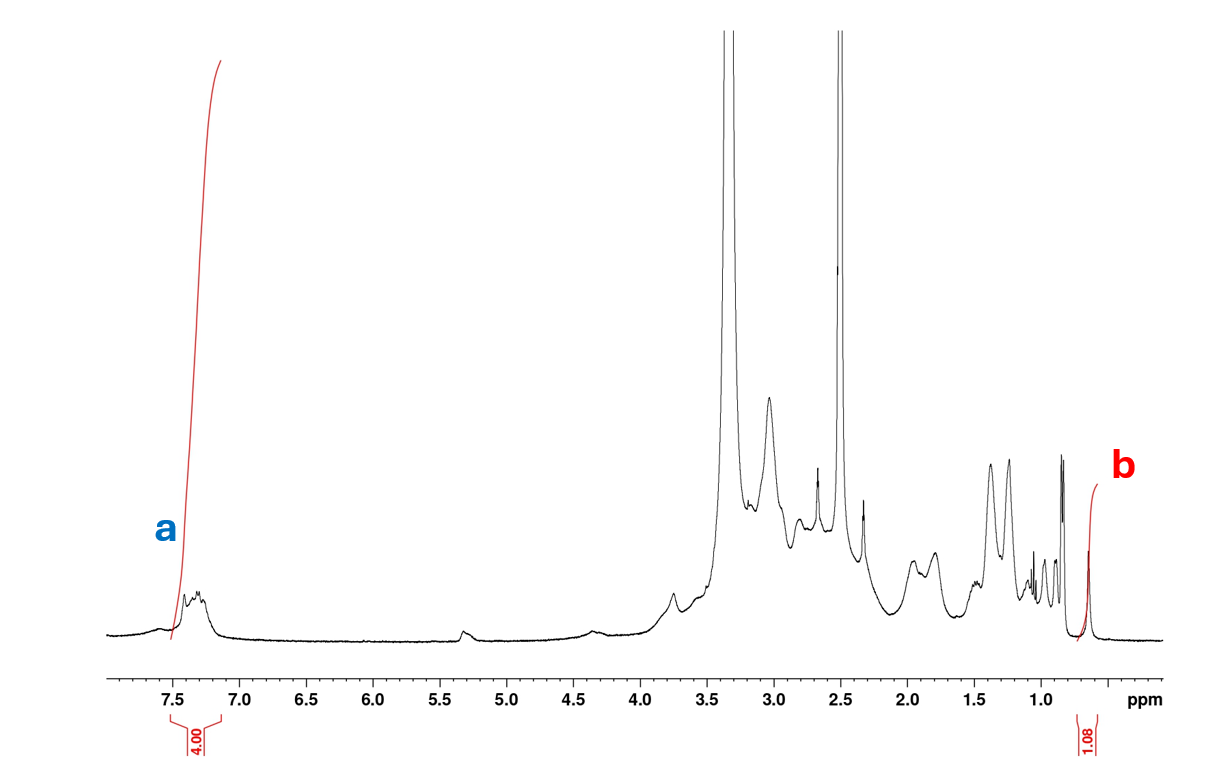

**Supplementary Figure 2.** Typical ^1^H-NMR spectrum of PAMD-C used in the determination of the cholesterol content ( spectrum of PAMD-C17 in DMSO-d6 is shown).

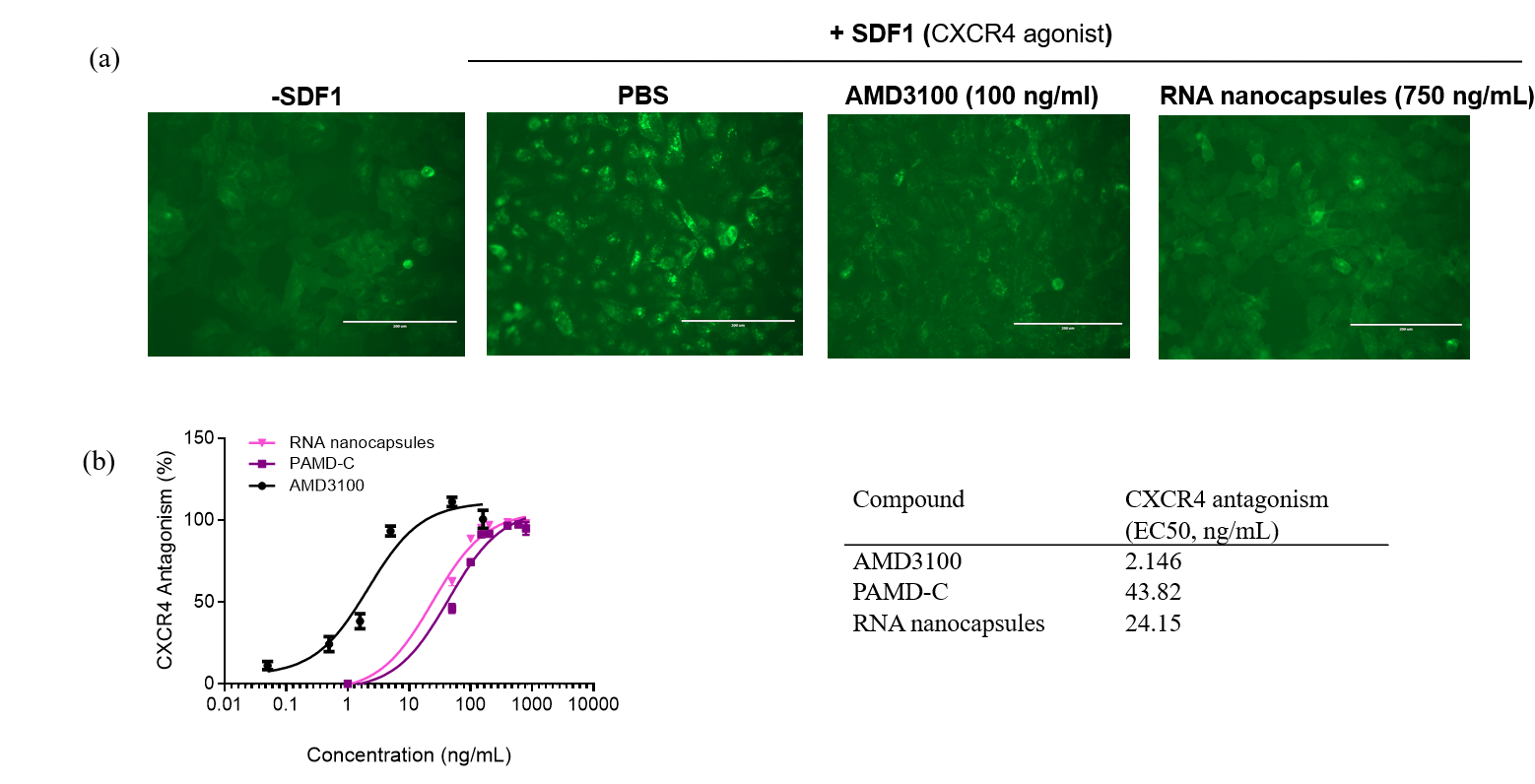

**Supplementary Figure 3. CXCR4 antagonism of RNA nanocapsules.** a) CXCR4 receptor redistribution assay in U2OS cells expressing GFP-tagged CXCR4(green). Scale bar = 100 um. b) EC_50_ values determined from the receptor redistribution assay (n=3). AMD3100 was used as the positive control.

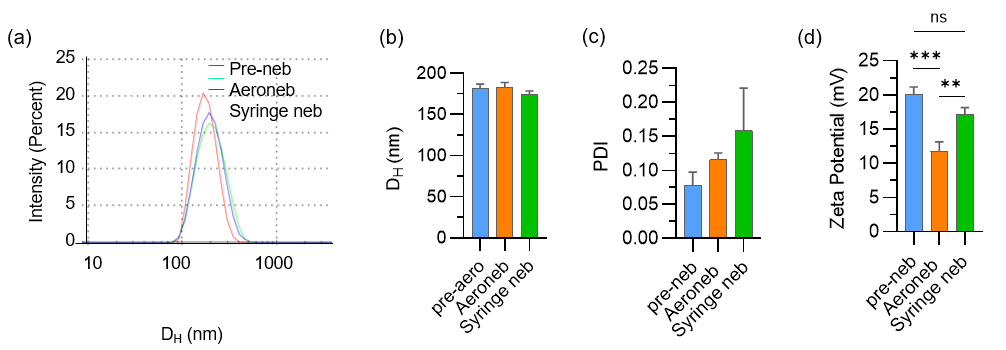

**Supplementary Figure 4. Formulation and characterization of neat and aerosolized PAMD-C@PFOB nanoemulsions.** a) Size distribution of nanoemulsion pre- and post-aerosolization. b) Mean hydrodynamic size, c) mean polydisperisty index (PDI), and d) mean zeta potential of nanoemulsions pre- and post-aerosolization by DLS.

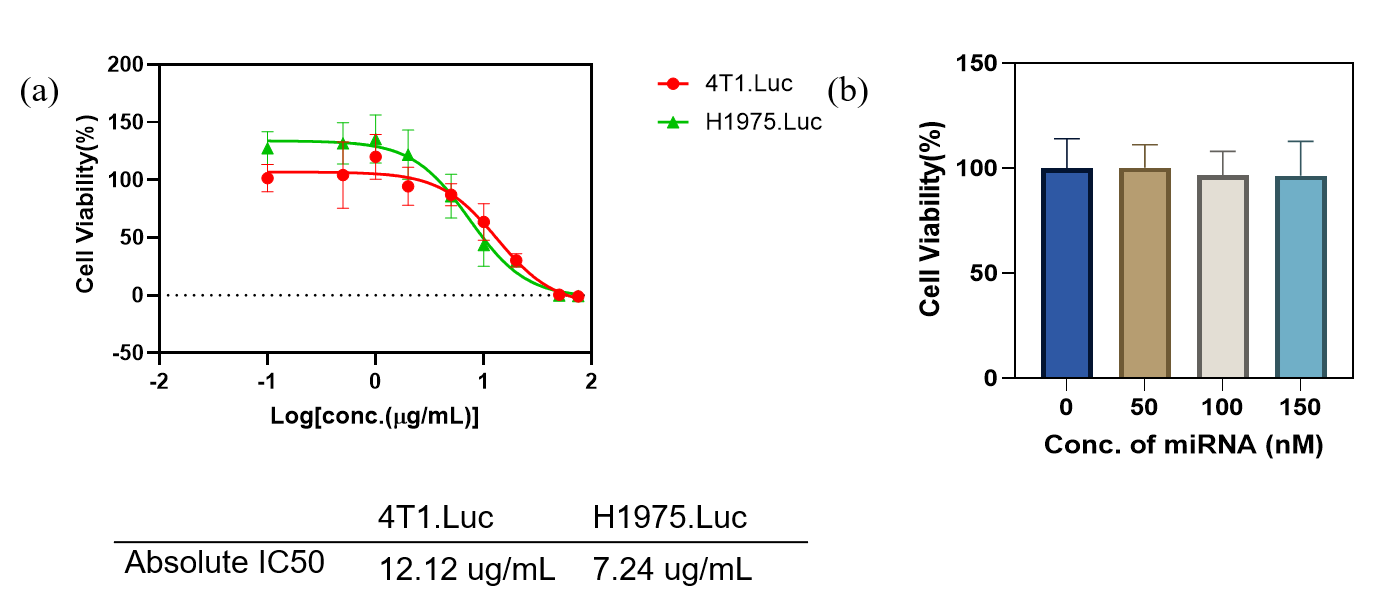

**Supplementary Figure 5. Cytotoxicity of PAMD-C@PFOB nanoemulsions and PAMD-C@PFOB/miRNA nanocapsules.** a) Absolute IC_50_ values of PAMD-Ch@PFOB in 4T1-Luc and NCI H1975-Luc cell lines. b) Cytotoxicity of RNA nanocapsules (w/w 4) at increasing concentration of miRNA in NCI-H1975-Luc cells.

**Supplementary Figure 6. Cellular uptake of PAMD-C@PFOB/miRNA nanocapsules.** a) Confocal microscopy images demonstrating cellular uptake of RNA nanocapsules (150 nM miRNA) after 6 h incubation in 4T1 cells. Scale bar = 20 μm. b) Cellular uptake of RNA nanocapsules at different w/w ratios by flow cytometry. n = 3 per group , *** p < 0.001.

| (a) Bare: PAMD-C + 1-fluorobutane  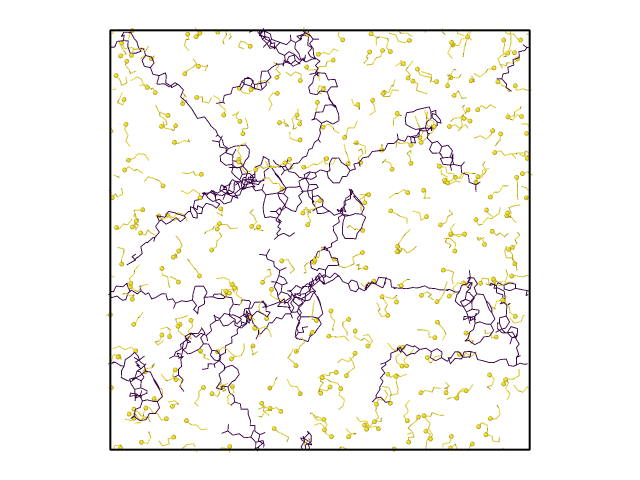  9 nm  9 nm | (b) |  | |
| --- | --- | --- | --- |
|  | 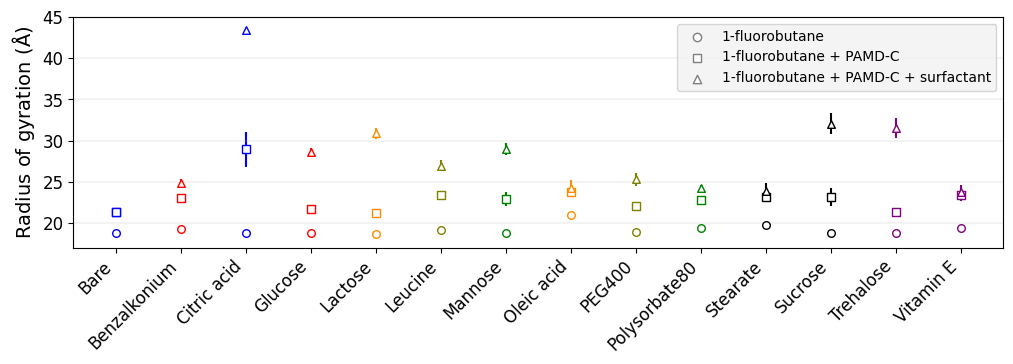 | | |
| (c)  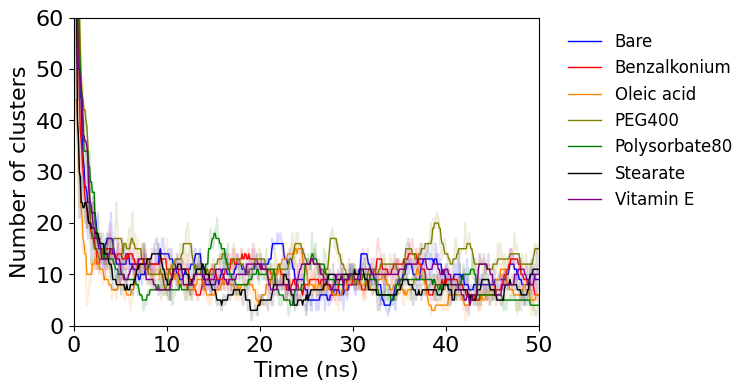 | | | (d)  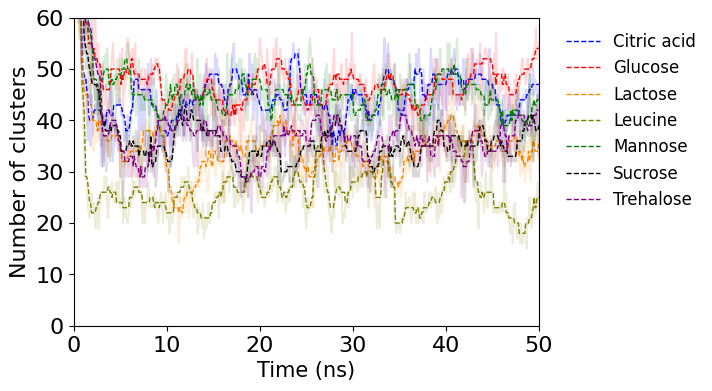 |
| (e) 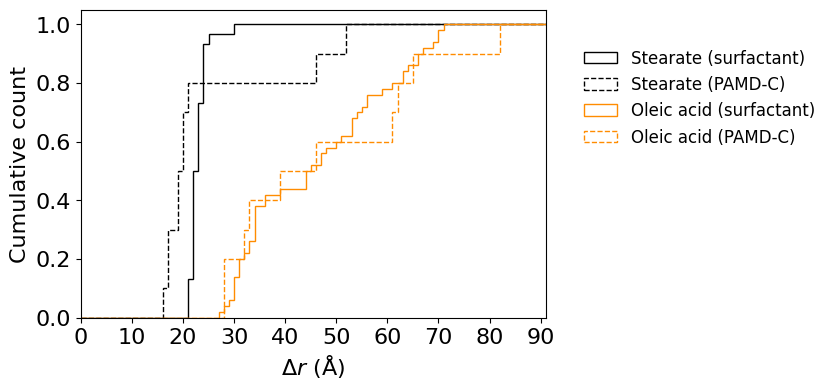 | | | (f)  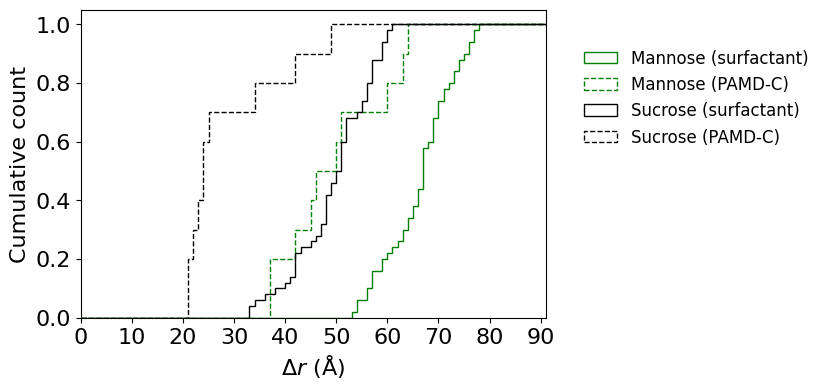 |
| **Supplementary Figure 7:** **AAMD simulations to study self-assembly of PAMD-C@1-fluorobutane and various excipients.** (a) Initial setup for the bare system containing PAMD-C and 1-fluorobutane. (b) Radius of gyration for all systems. There are three markers for each system representing the calculation of k using just the 1-fluorobutane molecules (○), 1-fluorobutane + PAMD-C molecules (□), and 1-fluorobutane + PAMD-C + excipient molecules (△). The vertical error bars denote the averaged value from 40-45 ns in the simulation. (c) and (d) show the evolution of number of clusters. (e) and (f) show the cumulative normalized histogram for $\Delta r$ (defined in the text). The distribution of $\Delta r$ is calculated between 40 ns and 45 ns (100 ps interval) and averaged to generate the plots. | | | |

As illustrated in Supplementary Figure 7a, the initial configuration was generated using a random dispersion of PAMD-C molecules and 1-fluobutane molecules in water. To examine the structural integrity of the particles, the radius of gyration ($k$) was calculated for sub-structures and is summarized in Supplementary Figure 7b. When only 1-fluorobutane molecules are included, the value of $k$ varies between 18-21 Å for all cases (bare system and the 13 excipients), indicating that the 1-fluorobutane molecules strongly interact with each other to form the core structure in all cases. When 1-fluorobutane and PAMD-C molecules are included*,* the value of $k$ increases for all systems, as expected from the snapshots. When 1-fluorobutane, PAMD-C, and excipient molecules are included, the change in $k$ can be used to infer the binding tendency of the excipient molecules. For benzalkonium, oleic acid, PEG400, polysorbate80, stearate, and vitamin E, the change in $k$ (difference between the △ and □ markers) is small, indicating that the excipient molecules are strongly bound to the particle. In contrast the large change in $k$ for citric acid, glucose, lactose, leucine, mannose, sucrose, and trehalose confirms that these molecules are randomly dispersed.

Further, clustering analysis presented in Supplementary Figure 7(c,d) shows the qualitative differences between the two sets of excipients. The number of clusters for excipients in Supplementary Figure 7c is comparable to the number of clusters in the bare system (10-15), implying that these excipient molecules are bound to the particle. On the other hand, number of clusters varies between 20-60 for the excipients in Supplementary Figure 7d. Inspecting the snapshots in Figure 2i (main text) shows that few 1-fluorobutane molecules are not part of the spherical core and are randomly dispersed. While these individual molecules contribute to the cluster count in Supplementary Figure 7(c,d) , they do not inhibit the tendency of the excipient to bind to the particle.

Lastly, Supplementary Figure 7(e,f) show the cumulative normalized histograms for $\Delta r$. For each PAMD-C or excipient molecule $i$,$\Delta r_{i}=\left\| r_{i}^{\mathrm{COM}}-r_{\mathrm{core}}^{\mathrm{COM}} \right\|$ is the Euclidean distance, where $r_{i}^{\mathrm{COM}}$ is the COM of the $i^{\mathrm{th}}$ molecule and$r_{\mathrm{core}}^{\mathrm{COM}}$ is the COM of all 1-fluorobutane molecules (which form the core) in the system. This calculation is done at each time $t$ in the simulation. Supplementary Figure 7e shows examples for stearate and oleic acid. In both cases, the dashed and solid histograms (calculation of $\Delta r$ for PAMD-C and excipient molecules respectively) are close to each other, which convey that the PAMD-C and excipient molecules are at comparable radial distances from $r_{\mathrm{core}}^{\mathrm{COM}}$. In other words, stearate and oleic acid bind strongly to the particle. On the other hand, the separation between the dashed and solid histograms for mannose and sucrose is much larger (Supplementary Figure 7f), and indicate the weak interactions between the excipients and the particle. Additionally, the snapshots for mannose and sucrose in Figure 2i (main text) show interactions between the particle and its periodic image. Such interactions are reduced for systems containing stearate, oleic acid, or other strong excipients, after stabilization of the particle (>30 ns).

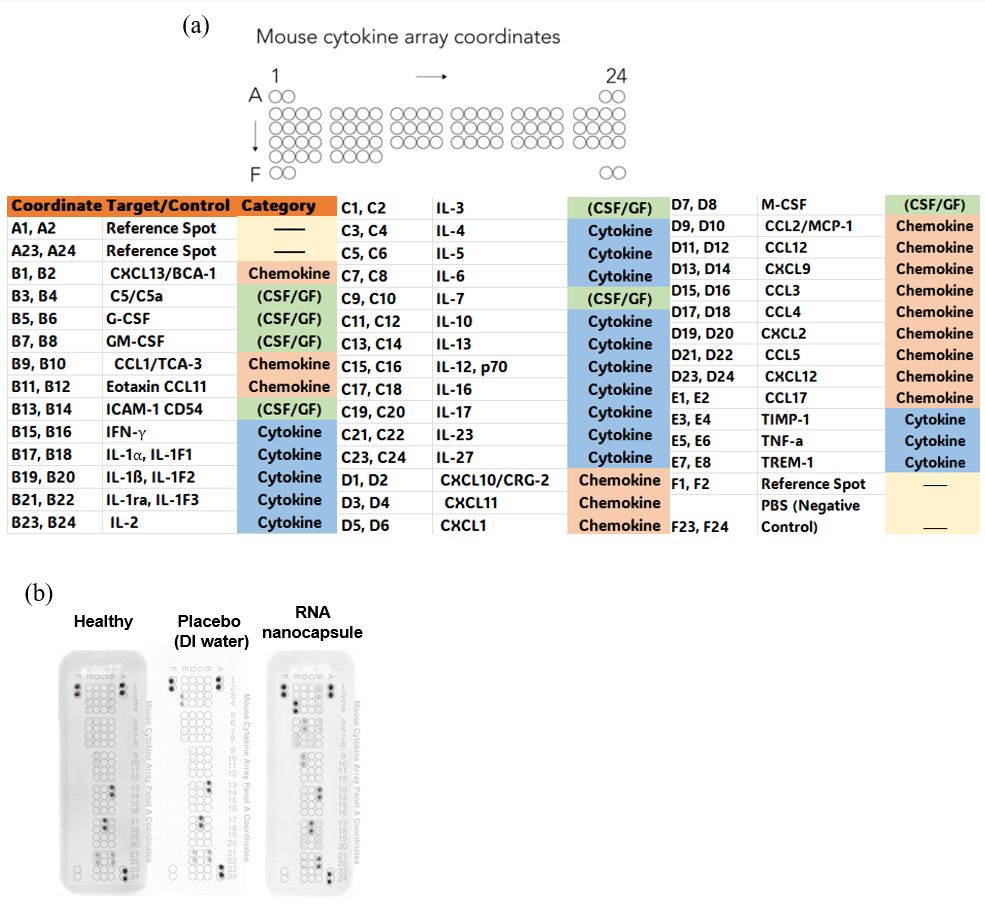

**Supplementary Figure 8. Serum cytokine, chemokine, and acute-phase protein expression after RNA nanocapsule administration.** a) Mouse cytokine array coordinates. b) Raw images of Proteome Profiler Mouse Cytokine Array showing differential expression of acute-phase proteins across groups, 12 h post- administration of RNA nanocapsules (60 µl, 20µg RNA) and DI water (60µl).

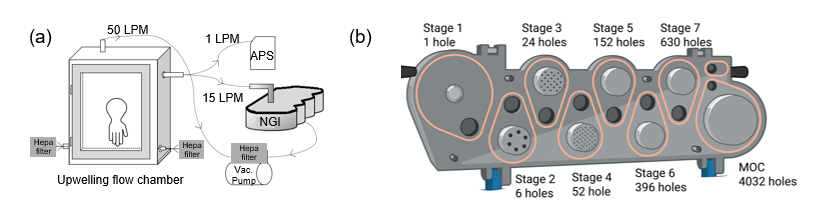

**Supplementary Figure 9. Characterization of PAMD-C@PFOB/miRNA nanocapsule aerosols.** (a) Schematic of upwelling flow chamber with NGI and APS units. (b) Schematic of the Next Generation Impactor (NGI) from MSP, a high-performance cascade impactor with seven stages and a Micro-Orifice Collector (MOC) at the end (*Figure created with Biorender.com)*

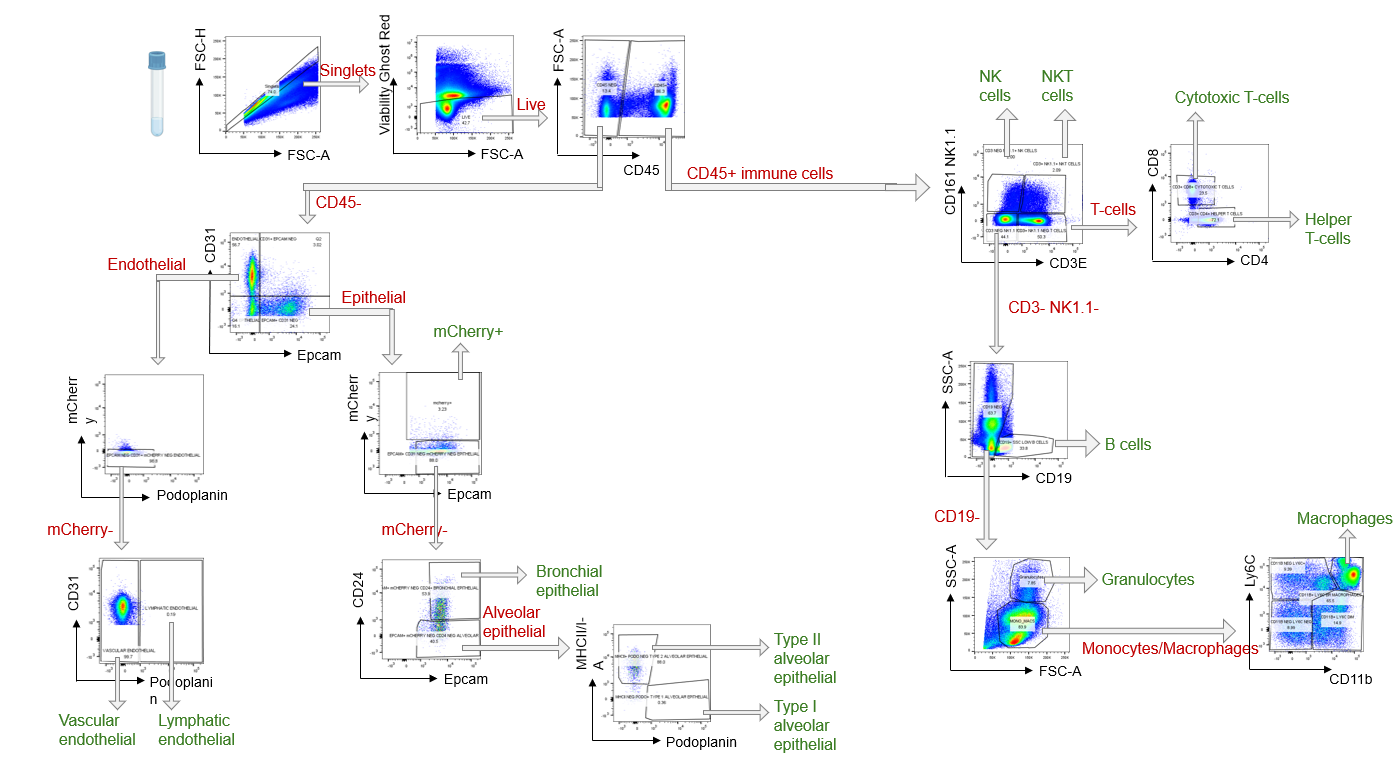

**Supplementary Figure 10.** Gating strategy for lung epithelial, endothelial, and immune cell populations used for assessing differential uptake of labelled miRNA.

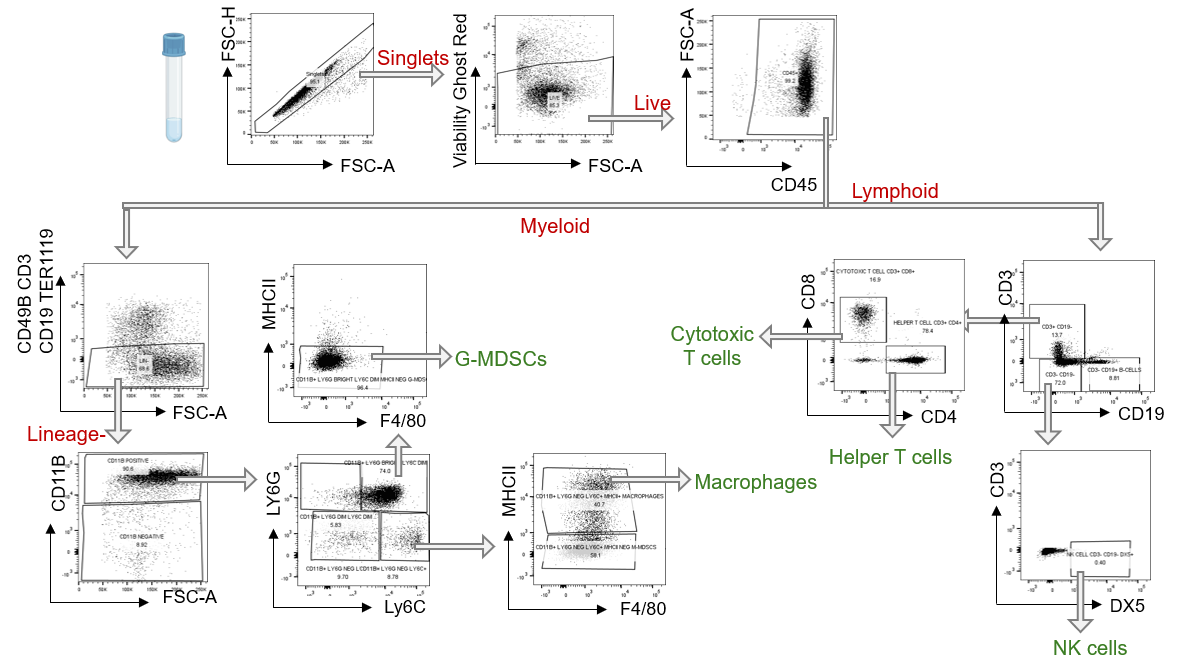

**Supplementary Figure 11.** Gating strategy for identification of distinct immune cell populations used for assessing immunomodulation in the tumor microenvironment in response to miR34a therapy.
